# SREBP-mediated gene expression regulation is essential for the intrinsic fungicide tolerance and antagonism in the fungal biocontrol agent *Clonostachys rosea*

**DOI:** 10.1101/2024.02.26.582079

**Authors:** Edoardo Piombo, Georgios Tzelepis, Alma Gustavsson Ruus, Vahideh Rafiei, Dan Funck Jensen, Magnus Karlsson, Mukesh Dubey

**Author notes:** Should be considered the joint first author (Edoardo Piombo, Georgios Tzelepis).

## Abstract

Sterol regulatory element-binding proteins (SREBPs) are a family of transcription factors known to regulate sterol biosynthesis and homeostasis in fungi. For this reason they have a role in several biological processes, including virulence, fungicide tolerance, hypoxia adaptation, lipid and carbohydrate metabolisms, and iron homeostasis. While the biological function of SREBPs in yeast and filamentous fungal species pathogenic to humans and plants is known, their role in fungal biocontrol agents (BCAs) is still elusive. This study aimed to investigate the biological and regulatory function of SREBPs in the BCA *Clonostachys rosea*, with a focus on their role in fungicide tolerance, hypoxia adaptation and antagonisms. The *C. rosea* genome contains two genes (*sre1* and *sre2*) coding for SREBPs and one gene each coding for Insulin induced gene (INSIG) and SREBP cleavage-activating protein (SCAP), required for SREBP-mediated ergosterol biosynthesis in fungi. Deletion of *sre1* resulted in mutants with pleiotropic effects, including the reduced ability to grow on media supplemented with proline (active ingredient prothioconazole) and cantus (active ingredient boscalid) fungicides, hypoxia mimicking agent CoCl2, cell wall stressor SDS, and increased growth rate on medium supplemented with caffeine, compared with *C. rosea* wild type (WT). In addition, the antagonistic ability against the fungal hosts *Botrytis cinerea* and *Rhizoctonia solani* was affected when *sre1* was deleted. However, no significant difference between *sre2* deletion strains and *C. rosea* WT was found for any of the tested phenotypes. To investigate the regulatory role of SRE1, the transcriptome of *C. rosea* WT and a *sre1* deletion strain was analyzed. The transcriptome analysis identified differentially expressed genes in the *sre1* deletion strain associated with carbohydrate and lipid metabolism, respiration, iron homeostasis, and xenobiotic tolerance. Moreover, genes coding for polyketide synthases and chitinases with a proven antimicrobial role were downregulated in the mutant, corroborating the reduced antagonism phenotypes. In summary, this work sheds light on the regulation role of transcription factor SRE1 while also exploring its effect on regulating the antagonistic activity and fungicide resistance of *C. rosea*, giving us helpful knowledge to design applications of this organism in IPM strategies.

## Introduction

Integrated Pest Management (IPM) is a holistic and sustainable approach to pest management that aims to minimise the use of chemical pesticides, thereby diminishing risks to human health and the environment (Deguine et al., 2021). Biological control, which is defined as the use of living organisms (biological control agents, BCAs) for controlling insect pests, weeds and plant diseases, is one of the essential components of IPM and is considered crucial to reducing the use of pesticides in future agriculture production systems (Stenberg et al., 2021). Due to the inherent ability of certain BCAs to tolerate a relatively higher dose of specific fungicides compared with doses suggested for controlling fungal plant pathogens (Chaparro et al., 2011; Dubey et al., 2016, 2014a; Jensen et al., 2011; Tzelepis and Lagopodi, 2011; Wedajo, 2015), they can be combined with a low or full dose of compatible pesticides either simultaneously or in rotation, resulting in an enhanced degree of disease suppression of crop plants (Ons et al., 2020; Thambugala et al., 2020). A combined application of fungal BCAs, for example *Clonostachys rosea* or *Trichoderma atroviride* with 1/10 of the field dose of triticonazole, showed disease control comparable to the full dose application of triticonazole on wheat caused by *Fusarium culmorum* (Roberti et al., 2006). Similarly, application of *Trichoderma* spp. with lower doses of various fungicides increased their efficacy in controlling *F. solani* and *F. oxysporum* in beans, *Cochliobolus heterostrophus* in corn and *Sclerotinia sclerotium* on oilseed rape (Abd-El-Khair et al., 2019; Wang et al., 2019; Hu et al., 2016). Although the combined application of BCAs with fungicides is considered a promising approach for future IPM strategies, a comprehensive knowledge of the inherent ability of BCAs to tolerate fungicides and their underlying mechanisms will help optimize the efficacy of BCA-fungicide combinations, which is of utmost importance for large-scale application under field conditions.

The development of fungicide tolerance is an evolutionary process. It can happen through various mechanisms, including alterations in cellular processes involved in fungicide uptake, target site modification and enforcement, metabolic detoxification and epigenetic changes altering gene expression (Yin et al., 2023). As the cell membrane is essential for fungal survival, many fungicides target fungal cell wall components, including sterol (Yin et al., 2023). Ergosterol is an integral part of the fungal cell membrane and crucial for fungal survival; thus, enzymatic steps of ergosterol biosynthesis are widely used as a drug and fungicide target for controlling fungal infections. The ergosterol biosynthesis is a complex process involving multiple enzymatic reactions and regulated by the sterol regulatory element-binding protein (SREBP) family of transcription factors. At optimum ergosterol concentration, SREBP is bound to the SCAP (SREBP cleavage activating protein), and the complex is retained at the endoplasmic reticulum (ER) by associating with INSIG (Yang et al., 2002). SCAP dissociates from INSIG at low ergosterol levels, releasing the SREBP-SCAP complex. The complex is then further transported to the Golgi apparatus, where the N-terminus of the SREBP is released from SCAP by proteolytic cleavage by two proteases (Espenshade and Hughes, 2007). The release enables SREBP to travel to the nucleus to induce gene expression of proteins involved in lipid biosynthesis and uptake (Espenshade and Hughes, 2007). The regulatory role of SREBPs is carried out by a basic helix loop helix (bHLH) leucine zipper DNA binding domain with a unique tyrosine residue at the N-terminus (Bien and Espenshade, 2010), while interaction between SREBP and SCAP is mediated by a domain of unknown function (DUF2014) (Maguire et al., 2014). In addition to sterol homeostasis and uptake, SREBPs regulate several biological processes in fungi, including carbohydrate metabolism, fungicide tolerance, hypoxia adaptation, and iron homeostasis (Espenshade and Hughes, 2007). The role of SREBPs in fungal virulence is also demonstrated (Espenshade and Hughes, 2007). However, while the role of SREBPs in all these processes has been shown in yeast and filamentous fungal species pathogenic to humans and plants, their role in fungal BCAs is still elusive. This study aimed to investigate the regulatory function of SREBPs in fungal BCAs, with a focus on exploring the mechanism by which they induce fungicide tolerance and their role in the regulation of biocontrol-related genes.

To achieve the aim we used *Clonostachys rosea*- a soil-borne filamentous fungus commonly known for its biocontrol capability against various plant-pathogenic fungi (Funck Jensen et al., 2021; Sun et al., 2020). Competition for nutrients and space, mycoparasitism, interference competition through antibiosis, plant root colonization and induction of plant defence responses are important biocontrol traits of *C. rosea* (Funck Jensen et al., 2021; Sun et al., 2020). Moreover, the tolerance of *C. rosea* to toxic fungal compounds has also been shown to contribute to its biocontrol ability (Dubey et al., 2016, 2014a; Kosawang et al., 2014). In addition, *C. rosea* has shown tolerance to chemical fungicides of various modes of action (Dubey et al., 2014a; Jensen et al., 2011; Macedo et al., 2012; Roberti et al., 2006; Tzelepis and Lagopodi, 2011) making it a promising *Candida*te for IPM strategy.

We hypothesized that SREBPs contribute to intrinsic fungicide tolerance and biocontrol traits in *C. rosea* by regulating ergosterol metabolism and xenobiotic efflux. Our results show that *C. rosea* contains genes required for SREBP-mediated ergosterol homeostasis. Furthermore, by generating gene deletion and complementation strains, we demonstrate the role of SREBPs in fungicide tolerance and antagonisms in *C. rosea*. Finally, through comparative transcriptome analysis of the SREBP deletion strain and *C. rosea* wildtype (WT), we elucidated the SREB-mediated transcriptional reprogramming of genes associated with lipid and carbohydrate metabolism, xenobiotic tolerance, and antagonism.

## Material and Methods

### Fungal strains and culture conditions

*C. rosea* strain IK726 wild type (WT) and mutants derived from it, *Botrytis cinerea* strain B05.10, *Fusarium graminearum* strain PH-1, *Rhizoctonia solani* SA1 were maintained on potato dextrose agar (PDA; Oxoid, Cambridge, UK) medium at 20°C. The yeast strains were maintained on YPD medium (in g/L; yeast extract 10, peptone 20, dextrose 20) at 30 °C. Unless otherwise specified, the Czapek-dox (CZ) medium (Sigma-Aldrich, St. Louis, MO) was used for phenotypic analyses.

### Gene identification and sequence analysis

The *C*. *rosea* strain IK726 genome version 1 (Karlsson et al., 2015) and version 2 (Broberg et al., 2018) were screened for genes encoding SREBP, INSIG and SCAP by BLASTP analysis. The presence of conserved domains was analysed with the Simple modular architecture research tool (SMART) (Letunic et al., 2009), InterProScan (Jones et al., 2014) and conserved domain search (CDS) (Marchler-Bauer et al., 2011). The presence of Tyrosine residues in SREBPs in specific spacing patterns was analysed manually. TOPCONS (http://topcons.net/; Tsirigos et al., 2015) and TMHMM-2.0 were used to predict the transmembrane domain and signal peptide prediction. Amino acid (aa) sequence alignment was performed using ClustalW2 (Larkin et al., 2007) with default settings for multiple sequence alignment.

### Comparative genomics analysis

All the published proteomes of Hypocreales species were downloaded from Mycocosm (Ahrendt et al., 2022). The genera *Fusarium*, *Metarhizium*, *Ophiocordyceps*, *Stachybotrys*, *Tolypocladium* and *Trichoderma* had a high number of annotated species, and therefore, only three species per genus were considered (**Supplementary table 1**). Each proteome was annotated with InterProScan v. 5.48-83.0 (with options “--iprlookup --goterms --pathways”) (Jones et al., 2014). Proteins showing both Pfam and InterProScan family “Insulin-induced protein” (PF07281 and IPR025929) were considered as putative INSIG proteins. Proteins showing family “Sterol regulatory element-binding protein cleavage-activating” (IPR030225) and the sterol sensing domain (IPR000731) were considered as putative SCAP proteins. Proteins having both domain bHLH (IPR011598) and domain “Sterol regulatory element-binding protein 1, C-terminal” (IPR019006) were considered to be putative SRE1 homologs. SRE2 proteins are short, and the only domain-based filtering in their prediction was the presence of the bHLH domain, resulting in a high number of *Candida*tes. For this reason, they were further filtered by excluding proteins with no blast matches on any of the SRE proteins predicted in the work of Chung et al. (2019). This last analysis was carried out with BLASTP v. 2.11.0 (with options “-e value 1e-10 -qcov_hsp_perc 80”) (Altschul et al., 1990). All the considered proteins are listed in **Supplementary table 1**.

### Domain-wise phylogenetic analysis

INSIG and SCAP proteins were identified as described previously in the species considered in the work by Chung et al. (2019), namely *Aspergillus fumigatus* (GCF_000002655.1), *Aspergillus nidulans* (GCF_000011425.1), *Candida albicans* (GCF_000182965.3), *Candida graminicola* (GCF_000149035.1), *F. graminearum* (GCF_000240135.3), *Fusarium oxysporum* (GCF_000271745.1), *Magnaporthe oryzae* (GCF_000002495.2), *Neurospora crassa* (GCF_000182925.2), *Saccharomyces cerevisiae* (NM_001184673.1) and *Schizosaccharomyces pombe* (NM_001018252.2). SRE proteins in these species were not identified through a new analysis, but the same proteins utilized in Chung et al. (2019) were considered.

One phylogenetic tree was obtained for each of the INSIG, SCAP and SREBP classes, considering only the domain PF07281 in INSIG proteins, domain IPR000731 in SCAP proteins and domain IPR011598 in SREBPs. Domain locations were predicted with InterProScan v. 5.48-83.0 (Jones et al., 2014), extracted with SAMtools v. 1.11 (Danecek et al., 2021) and aligned with MAFFT v. 7.453 (with options “--maxiterate 1000 –localpair”) (Katoh and Standley, 2013). The trees were obtained with IQ-TREE v. 2.1.3 (with options “- m MFP -b 1000 -T 1 -safe”) (Minh et al., 2020). The best-fit models chosen by the ModelFinder function of IQ-TREE were Q.plant+G4 for INSIG proteins, mtZOA+G4 for SCAP proteins and JTT+G4 for SRE proteins. After 1000 bootstraps, the SRE tree nodes with bootstrap values less than 50% were condensed through MEGA v. 10.0.5 to improve readability (Kumar et al., 2018), and the trees figures were improved with Scientific Inkscape (https://github.com/burghoff/Scientific-Inkscape/tree/main).

### Yeast-two-hybrid assays

For the yeast-two-hybrid assays, SRE1, SRE2, and SCAP genes were amplified from *C. rosea* cDNA using gene-specific primers (**Supplementary table 2)** and subcloned to the pDONOR/Zeo donor vector (Thermo Fisher, MA). The donor vectors were cloned either to the pGADT7-GW prey or the pGBKT7-GW bait plasmids (Takara, Kusatsu, Japan) and simultaneously transformed to the *Saccharomyces cerevisiae* AH109 strain (Clontech, CA) as described above. Transformation with empty vectors was used as negative control. Positive colonies were selected on Synthetic minimal (SD) -Leu, -Trp, media and potential protein interactions were evaluated on SD -His, -Ade, -Leu, -Trp media. Five replicates per interaction have been used.

### Heterologous expression of *sre1* and *sre2* in *Saccharomyces cerevisiae*

For heterologous expression in *S. cerevisiae*, the *sre1* and *sre2* genes were amplified using cDNA from *C. rosea* and cloned to the pYES-2 vector driven by *GAL1* promoter, using the GeneArt™ Seamless Cloning and Assembly Enzyme Mix (Thermo Fisher Scientific, MA) according to manufacturers’ instructions. The vectors were transformed using a polyethene glycol-based protocol in the BY4742 strain (Agatep et al., 1998) and positive transformants were selected on Synthetic Complete (SC) -Ura media. Transformation with the empty pYES-2 vector was used as a negative control. For gene induction, overexpression strains were precultured in SC -Ura media with 1% raffinose to reach the log phase. Then, the OD_600_ was adjusted to 0.3 and transferred to SC -Ura media supplemented with 2% galactose. The growth of *S. cerevisiae* strains was investigated in SC -Ura medium supplemented with 1.5 ppm prothioconazole dissolved in 50 % DMSO (Merck, NJ) and measuring the OD_600_ in SpectraMax Gemini™ XPS/EM microplate reader (Molecular Devices, CA) at 30 °C in a time-course assay. In control treatment, prothioconazole was replaced with an equal volume of 50% DMSO. The experiment was performed with six biological replicates. The optimum concentration of prothioconazole was selected by successive screening of the *S. cerevisiae* strain to a prothioconazole concentration ranging from 0.1 ppm to 5 ppm.

### Construction of deletion vector, transformation, and mutant validation

The ∼ 1 kb 5’-flank and 3’-flank regions of *sre1* and *sre2* were amplified from the genomic DNA of *C*. *rosea* using gene-specific primer pairs as indicated in **Supplementary figure 1**. Gateway entry clones of the purified 5’-flank and 3’-flank PCR fragments were generated as described by the manufacturer (Invitrogen, Carlsbad, CA). The hygromycin resistance cassette (hygB) generated during our previous studies (Dubey et al., 2012) from the pCT74 vector, as well as a geneticin resistance cassette generated as a PCR product from the pUG6 vector (Güldener et al., 1996), were used. The gateway LR recombination reaction was performed using the entry plasmid of respective fragments and destination vector pPm43GW (Karimi et al., 2005) to generate the deletion vectors. A complementation cassette for *sre1* was constructed by amplifying the full-length sequence of *sre1,* including more than 1 kb upstream and around 500 bp downstream regions from the genomic DNA of *C. rosea* WT (**Supplementary table 2; Supplementary figure 1).** The amplified DNA fragments were purified and integrated into the destination vector pPm43GW using Gateway cloning technology (Invitrogen, CA) to generate complementation vectors.

*Agrobacterium tumefaciens*-mediated transformation (ATMT) was performed based on a previous protocol for *C. rosea* (Utermark and Karlovsky, 2008). Transformed strains were selected on plates containing hygromycin for gene deletion and geneticin for complementation. Validation of homologous integration of the deletion cassettes in putative transformants was performed using a PCR screening approach with primer combinations targeting the hygB cassette and sequences flanking the deletion cassettes **(Supplementary figure 1**), as described previously (Dubey et al., 2013a, 2013b). The PCR-positive transformants were purified by two rounds of single spore isolation (Dubey et al., 2012). The transcript levels of *sre1* and *sre2* on WT, and respective gene deletion and complementation strains, were determined by RT-PCR using RevertAid premium reverse transcriptase (Fermentas, St. Leon-Rot, Germany) and their respective primer pairs (**Supplementary table 2; Supplementary figure 1**).

### Phenotypic analyses

Phenotypic analysis experiments were performed with *C*. *rosea* WT, three independent single deletion strains of *sre1* (Δ*sre1_1*, Δ*sre1_5*, Δ*sre1_15*) and *sre2* (Δ*sre2_14*, Δ*sre2_55*, Δ*sre2_104*) and *sre1* complemented strain Δ*sre1*+. Each experiment included three to five biological replicates (depending on the phenotype), and each experiment was repeated two times with similar results unless otherwise specified.

For growth rate analysis, a 3 mm diameter agar plug from the growing mycelial front was transferred to solid CZ medium or CZ medium containing fungicides proline EC 250 (Bayer Crop Science; active ingredient prothioconazole, azole group of fungicides, 0.0125 μg /ml), Cantus WDG (BASF Canada Inc; active ingredient boscalid, anilid group of fungicides, boscalid, 2000 μg/ml), Chipco green 75WG (Bayer Crop Science; active ingredient iprodione, dicarboximide group of fungicides, iprodione, 250 μg/ml), or Teldor WG 50 (Bayer Crop Science; active ingredient Fenhexamid; Hydroxyanilide group of fungicides, fenhexamid, 7500 μg/ml); hypoxia inducing agent cobalt chloride (Merck, NJ; CoCl_2_ 2.5 mM); cell wall stressors SDS (0.05%) or caffeine (0.1%). The optimum concentration of proline and CoCl_2_ was selected based on the screening of *C*. *rosea* to these compounds on CZ plates. The concentration of other fungicides and cell wall stress inducers used in this study is described in our previously published results (Dubey et al., 2014a, 2016). Colony diameter was measured four days post inoculation (dpi) at 20 °C. Agar plugs of *C. rosea* strains were inoculated on microscope slides with CZ medium supplemented with prothioconazole for microscopy observation of colony morphology. The mycelial edge of the colonies was photographed using a Leica DM5500M Microscope equipped with a Leica DFC360FX digital camera (Wetzlar, Germany).

Antagonistic behaviour against the phytopathogenic fungi (mycohosts), *B*. *cinerea, F. graminearum* and *R*. *solani* was tested using an *in vitro* plate confrontation assay on PDA medium as described before (Dubey et al., 2014a, 2014b, 2016). The growth of *B*. *cinerea, F. graminearum* and *R*. *solani* was measured daily until their mycelial front touched the *C*. *rosea* mycelial front. The growth of *C. rosea* strains over the mycohosts was measured until the fungus reached another side of the plate. The plate confrontation assay was performed in four biological replicates.

The biocontrol ability of the *C. rosea* strains against *F*. *graminearum* was evaluated using a fusarium foot rot assay (Dubey et al., 2020; Knudsen et al., 1995). In brief, surface sterilised wheat seeds were treated with *C. rosea* conidia (1e+07 conidia/ml) in sterile water, sown in moistened sand, and kept in a growth chamber after pathogen inoculation (Dubey et al., 2016, 2014b). Plants were harvested three weeks post-inoculation, and disease symptoms were scored on a 0-4 scale, as described before (Dubey et al., 2014b; Knudsen et al., 1995). The experiment was performed in five biological replicates, with 15 plants in each replicate.

### RNA sequencing

The transcriptome of *C. rosea* WT and Δ*sre1* strains was analysed in submerged liquid CZ medium and CZ medium amended with prothioconazole. Conidia of *C. rosea* strains were pre-cultivated for two days in 100 ml flasks containing 20 ml liquid CZ medium on a rotary shaker (100 rpm) at 25 °C after which the growth medium was amended directly with 0.03 ppm prothioconazole. After 24 h of incubation, fungal mycelia were harvested, washed in distilled water, frozen in liquid nitrogen and stored at -80 °C. The experiment was performed with four biological replicates. RNA extraction was done using the Qiagen RNeasy kit following the manufacturer’s protocol (Qiagen, Hilden, Germany). After DNaseI (Fermentas, St. Leon-Rot, Germany) treatment, the RNA quality was analysed using a 2100 Bioanalyzer Instrument (Agilent Technologies, Santa Clara, CA), and concentration was measured using a Qubit fluorometer (Life Technologies, Carlsbad, CA). For mRNA sequencing, the total RNA was sent for library preparation and paired-end sequencing at the National Genomics

Infrastructure (NGI), Uppsala, Sweden. Sequencing libraries were prepared using the TruSeq stranded mRNA library preparation kit (Illumina Inc. San Diego, CA), including polyA selection according to the manufacturer’s protocol (Illumina Inc. San Diego, CA). The mRNA libraries were sequenced on one NovaSeq SP flowcell with a 2×150 setup using the Illumina NovaSeq6000 system at the SNP&SEQ Technology Platform, Uppsala, Sweden.

For RNAseq analysis, the raw reads underwent adapter removal and quality trimming with the BBDuk tool from BBmap v. 38.9 (with options “ktrim=r k=23 mink=11 hdist=1 tpe tbo qtrim=r trimq=10 maq=10”) (Bushnell, 2019), and quality was checked with FastQC v. 0.11.9 (Andrews, 2010). The clean reads were mapped to the genome of *C. rosea* IK726 (GCA_902827195.2) using STAR v. 2.7.9a with default parameters (Dobin et al., 2013). The number of reads mapping to each gene was counted using featureCounts v. 2.0.1 (with options “-p -O --fraction -g Parent -t exon -s 2”) (Liao et al., 2014). The differentially expressed genes were then determined using the R package DESeq2 (Love et al., 2014) with a maximum FDR-adjusted p-value of 0.05 and a threshold on absolute log2(FC) of 1.

Gene ontology enrichment was performed by performing one-tailed Fisher’exact tests with an FDR threshold of 0.05 using BLAST2GO v. 5.2.5 (Conesa et al., 2005). The annotation of the *C. rosea* genome used for the analysis was the same one described in Piombo et al. (2021). Furthermore, differentially expressed genes involved in respiration, iron ion binding, sterol metabolism, and cytochrome P450 genes were predicted using the annotation of Piombo et al. (2021). ABC and MFS transporters were also identified, and they were assigned a class depending on the prediction in Broberg et al. (2021). The proteins encoded by these genes of interest were compared with the fungal section of the NCBI non-redundant database using BLAST (Altschul et al., 1990).

### Statistical analysis

Analysis of variance (ANOVA) was performed on gene expression and phenotype data using a General Linear Model approach implemented in Statistica version 13 (TIBCO Software Inc., Palo Alto, CA). Pairwise comparisons were made using the Fisher’s or Tukey-Kramer method at the 95% significance level. The students’ *t-test* was also performed on gene expression data.

## Results

### Identification and sequence analysis of SREBPs, INSIG and SCAP in *C. rosea*

Analysis of the *C*. *rosea* strain IK726 genome (Broberg et al., 2018; Karlsson et al., 2015) identified two genes predicted to encode SREBPs (CRV2T00000933_1, named *sre1*; CRV2T00003323_1, named *sre2*), one gene each encoding for INSIG (CRV2T00010728_1, named *insig*), and SCAP (CRV2T00000306_1, named *scap*). SRE1 was composed of 999 aa that contained a predicted basic Helix-Loop-Helix (bHLH) domain (PF00010, IPR011598) with a tyrosine (Y) residue unique to SREBP at the N-terminus and a domain of unknown function DUF2014 (PF09427) at C terminus. Like *S. pombe* Sre1+ and Sre2+ (Osborne and Espenshade, 2009), *C. rosea* SRE1 contains two transmembrane helices. SRE2 has a shorter protein sequence than SRE1, consisting of 328 residues and lacks DUF2014 domain and transmembrane helices (**Supplementary figures 2 and 3**). The characteristics of these proteins are presented in **Table 1**. Like previously characterized SRE sequences, the bHLH domain of *C. rosea* SRE proteins contains conserved HNxxExxYR and KxxxLxxAxxYxxxL motifs in the basic and helix regions. In contrast, the loop region is highly variable (**Figure 1**). A phylogenetic analysis using the bHLH domain of selected SREBP proteins (Chung et al., 2019) showed that *C. rosea* SRE1 belongs to Sordariomycetes SRE clade A (SRE clade one), while SRE2 is external to both clade one and clade two (Ruan et al., 2019). The phylogenetic tree obtained by comparing the *C. rosea* SCAP and INSIG proteins to the sequences used in Chung et al. (2019) showed that the *C. rosea* proteins clustered according to evolutionary distance (**Supplementary figure 3**).

**Figure 1:**
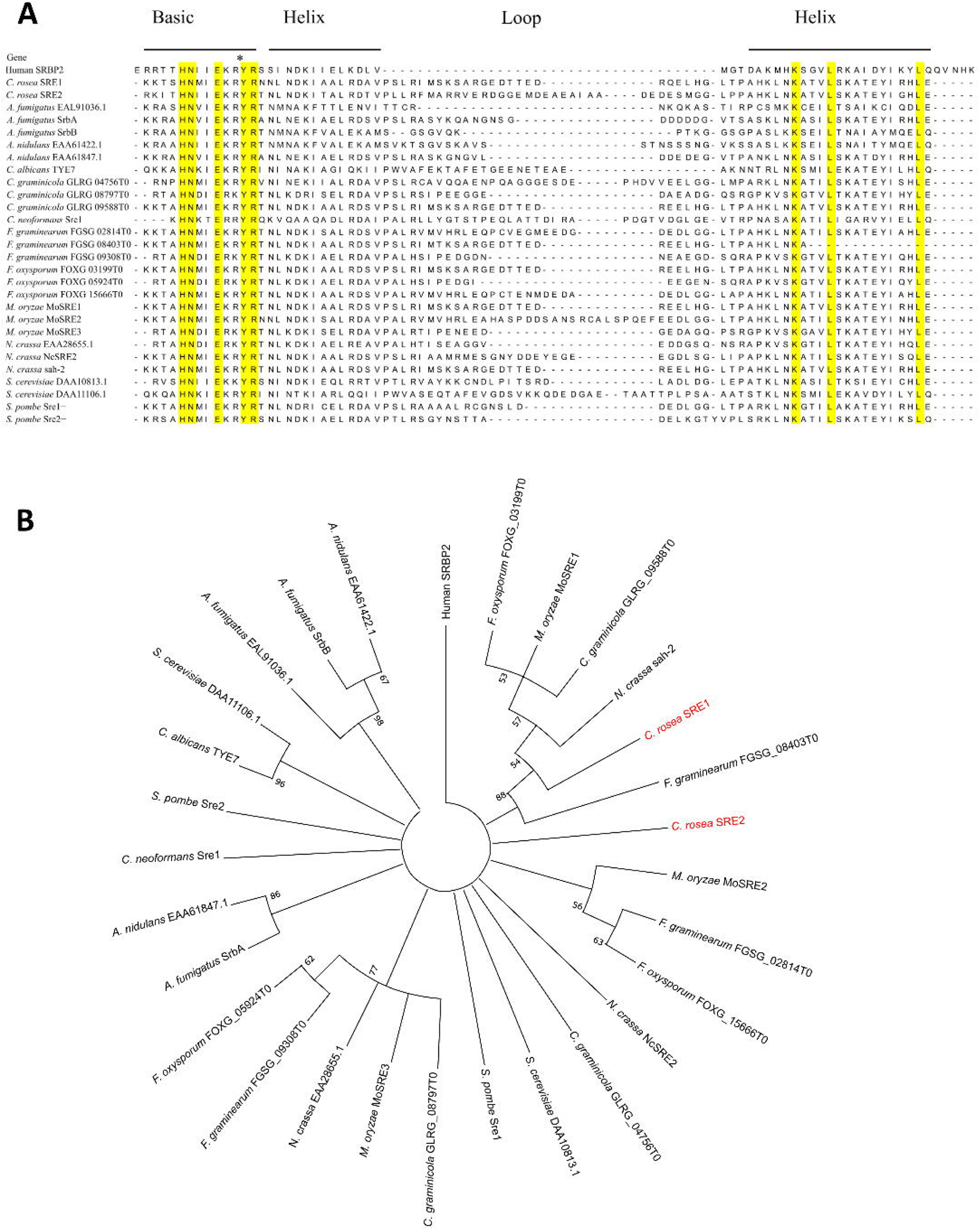
Alignment of the BHLH domain of SRE proteins (**A**) and phylogenetic tree based upon the same sequences (**B**). Besides *C. rosea* SRE1 and SRE2, the included proteins are the ones considered in the work of Chung et al. (2019).

**Table 1:** Characteristics of putative SREBPs, INSIG and SCAP in *C. rosea*.

| Name | Transcript ID | Length of genomic sequence | No of introns | Protein length | Transmembrane helices | Domain |
| --- | --- | --- | --- | --- | --- | --- |
| SRE1 | CRV2T00000933 | 3054 | 1 | 999 aa | 2 | DUF2014 (239 aa – 313 aa)<br>bHLH (581 aa – 845 aa) |
| SRE2 | CRV2T00003323 | 984 | 0 | 328 aa | 0 | bHLH (210 aa – 316 aa) |
| INSIG | CRV2T00010728 | 1292 | 2 | 407 aa | 6 | INSIG superfamily (165 aa – 392 aa) |
| SCAP | CRV2T000003061 | 3563 | 2 | 1127 aa | 8 | SCAP family (7 aa – 699 aa)<br>WD40 (554 aa – 563 aa;<br>606 aa – 640 aa);<br>SSD (318 – 457) |

The putative INSIG homolog (CRV2T00010728_1) in *C. rosea* is predicted to consist of 407 residues and contains the INSIG superfamily domain (PF07281, IPR025929) at aa position 165-392 and six transmembrane helices between aa position 160 and 388. The *C. rosea* SCAP is composed of 1127 aa with a SCAP family domain (IPR030225) consisting of sterol sensing domain (SSD; IPR000731, PF12349), nine transmembrane helices spanning between 36 aa – 936 aa, and two WD40 repeat motif (IPR001680) between 554 aa – 593 aa and 659 aa – 730 aa, mediating protein-protein interactions (**Table 1; Supplementary figure 2**).

To investigate the distribution of SREBP, INSIG and SCAP genes among fungal species from order Hypocreales, we searched 40 fungal genomes representing plant-pathogenic, insect-pathogenic, mycoparasitic and nematode-pathogenic lifestyles. SRE1 homologues were identified in all analysed species except insect pathogens *Escovopsis weberi* and *Cordyceps militaris*. At the same time, the gene coding for SRE2 is variably distributed, from one gene among the fungi with mycoparasitic and nematophagous lifestyles to three genes in plant pathogenic lifestyles such as *Claviceps purpurea*. Interestingly, the SRE2 gene is absent in specific insect pathogenic fungi (for example *Beauveria bassiana* and *Cordyceps militaris*) that belong to the family Cordycipitaceae, but other insect pathogens such as *Metarhizium anisopliae* and *M. anisopliae* from Clavicipitaceae family contain two SRE2 genes (**Supplementary table 3).** The gene copy number of genes coding for SCAP and INSIG are conserved among the species analysed in this study except for *Lecanicillium lecanii* and *Torrubiella hemipterigena*, which lack a SCAP homologue, and *Stachybotrys chartarum* lacking INSIG.

### SRE1, but not SRE2, interacts with SCAP

To investigate the physical interaction between SREBP and SCAP, we used Y2H, a well-established method to study protein-protein interactions (Brückner et al., 2009). Yeast strains co-transformed with SRE1 and SCAP were able to grow on the auxotrophic selection plates (-His, -Ade, -Leu, -Trp), while yeast strains co-transformed with SRE2 and SCAP failed to grow on selection plates. The results from the Y2H assay suggested a physical interaction between SRE1 and SCAP1 but not between SRE2 and SCAP (**Figure 2A).**

**Figure 2:**
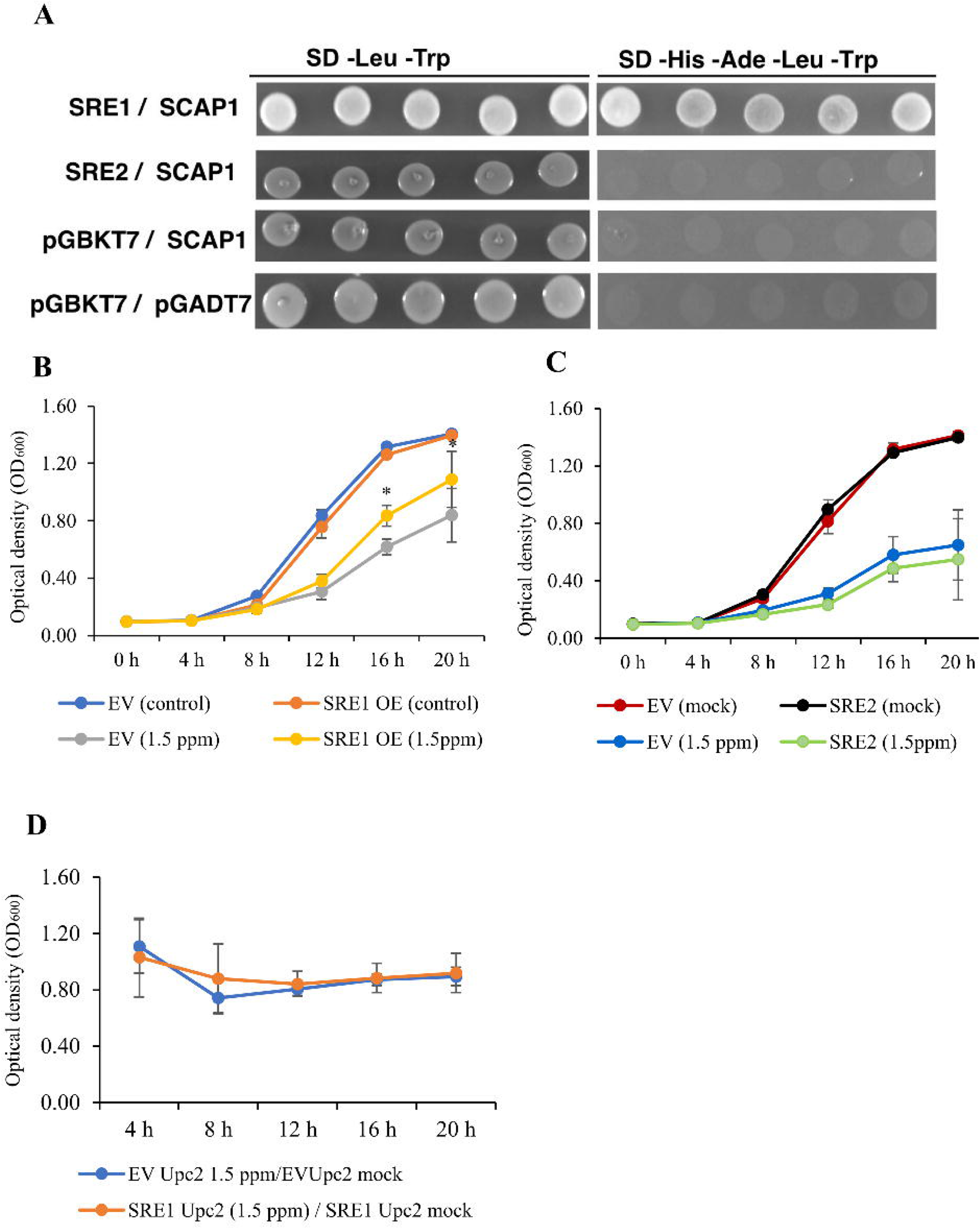
Analysis of the *sre1* and *sre2* genes in *Saccharomyces cerevisiae*. **A**: Yeast-two-hybrid assay between SRE1 or SRE2 (used as a bait in pGBKT7 vector) and SCAP1 (used as a prey in pGADT7 vector). Growth of yeast cells on SD-4 (-His, -Ade, -Leu, -Trp) selective media represents protein–protein interaction and growth on SD-2 (-Leu, -Trp) media confirms yeast transformation. Yeast transformed with the empty vectors were used as negative controls. **B**: Growth of *S. cerevisiae* overexpressing *sre1* upon exposure to 1.5 ppm prothioconazole. **C**: Growth of *S. cerevisiae* overexpressing *sre2* upon exposure to 1.5 ppm prothioconazole. **D**: Growth of *S. cerevisiae Upc2* mutants complemented with either *sre1* or *sre2* upon exposure to 1.5 ppm prothioconazole. In all assays, *S. cerevisiae* strains transformed with the empty pYES-2 vector (EV) and grown in equal volume of 50% DMSO were used as controls. Asterisks represent statistical significant differences based on t test (p < 0.05). Error bars represent standard deviation based on six biological replicates.

Overexpression of *C. rosea sre1* in *Saccharomyces cerevisiae* enhanced tolerance to prothioconazole.

To corroborate that *C. rosea* SRE1 and SRE2 are SREBPs, we generated *S. cerevisiae* strains overexpressing *C. rosea sre1* (SRE1 OE) and *sre2* (SRE2 OE). *S. cerevisiae* strain transformed with empty vector (EV) was used as control. The *S. cerevisiae* genome lacks SREBP, and sterol synthesis is regulated by a typical Gal4-type zinc finger transcription factor named UPC2, which differs from SREBP in protein structure. The tolerance of SRE1 OE and SRE2 OE to prothioconazole was determined by measuring their growth rates in yeast extract peptone dextrose (YPD) medium supplemented with 1.5 ppm prothioconazole. A significant 24% increase in growth of the *S. cerevisiae* SRE1 OE strain was found in the YPD medium supplemented with prothioconazole compared to the EV control at 12 hpi (*P* = 0.022), followed by 35 % and 30 % (*P* = 0.001) reduction at 16 hpi and 20 hpi, respectively (**Figure 2B**). In contrast, the *S. cerevisiae* SRE2 OE strain showed no significant difference in growth compared to the EV control at the tested time points (**Figure 2C**). No significant differences (*P* = 0.061) in growth between the EV strains and SRE1 OE or SRE2 OE strains in the absence of the prothioconazole showed the insertion of EV did not affect the growth of *S. cerevisiae,* and the observed phenotype is due to the expression of SRE1 (**Figure 2C**).

Since *C. rosea* SRE1 could enhance the tolerance of *S. cerevisiae* to prothioconazole, we overexpressed this gene in *S. cerevisiae Upc2* knock out strain (Δ*Upc2*) background (SRE1 Upc2) to investigate whether SRE1 is a functional analogue to Upc2. Since *S. cerevisiae* SRE1 Upc2 strain showed a reduced growth rate (P = 0.001) compared to that of the empty vector (EV Upc2) control (**Supplementary figure 4**), the growth rate of SRE1 Upc2 and EV Upc2 in the YPD medium with prothioconazole was normalized to the growth rate in the YPD medium. The result showed no significant differences (*P* ≥ 0.21) in the normalized growth rate between the SRE1 Upc2 strain and empty EV Upc2 (**Figure 2D**).

### Generation of *sre* deletion and complementation strains

To characterize the biological role of SRE1 and SRE2 in *C*. *rosea*, *sre1* and *sre2* deletion strains were generated by replacing the genes with the hygromycin resistance gene cassette hygB through ATMT **(Supplementary figure 1**). Successful gene replacement in hygromycin-resistant transformants was confirmed by PCR using primers following the procedure described previously (Dubey et al., 2012). The expected size of PCR fragments was amplified in Δ*sre1* and Δ*sre2* strains, while no amplification was observed in the WT (**Supplementary figure 1**). Furthermore, RT-PCR experiments using primers specific to the *sre1* and *sre2* sequences demonstrated the complete loss of *sre1* and *sre2* transcripts in each mutant, while expression of *sre1* and *sre2* was detected in the WT (**Supplementary figure 1**). The Δ*sre1* strain was complemented with *sre1*. Successful integration of the complementation cassette in mitotically stable mutants was confirmed by PCR amplification of geneticin-resistant selection cassette (**Supplementary figure 1**). RT-PCR from randomly selected geneticin positive Δ*sre1* strains using *sre1*-specific primer pairs demonstrated restored *sre1* transcription in Δ*sre1* complemented (Δ*sre1*+) strains. At the same time, no transcripts were detected in the parental deletion strains (**Supplementary figure 1**).

### Deletion of *sre1* affects *C. rosea* tolerance to fungicides and hypoxia

No differences in growth rate were detected between the deletion and WT strains on PDA or CZ media. The role of SRE in fungicide tolerance was investigated by comparing the growth rate of SRE deletion and WT strains on a CZ medium amended with proline, Teldor, Cantus or Chipkogreen. The growth rate of the Δ*sre1* strains was 51% and 22% lower (*P* < 0.001) than the WT growth rate on CZ amended with proline and Cantus fungicides, respectively. At the same time, no significant difference in growth rate was found between the deletion and WT strains on CZ amended with Chipkogreen or Teldor (**Figure 3A**). Microscopic observation of *C. rosea* colonies showed a deformed mycelial front of Δ*sre1* strain on PDA supplemented with proline fungicide (**Supplementary figure 5**).

**Figure 3:**
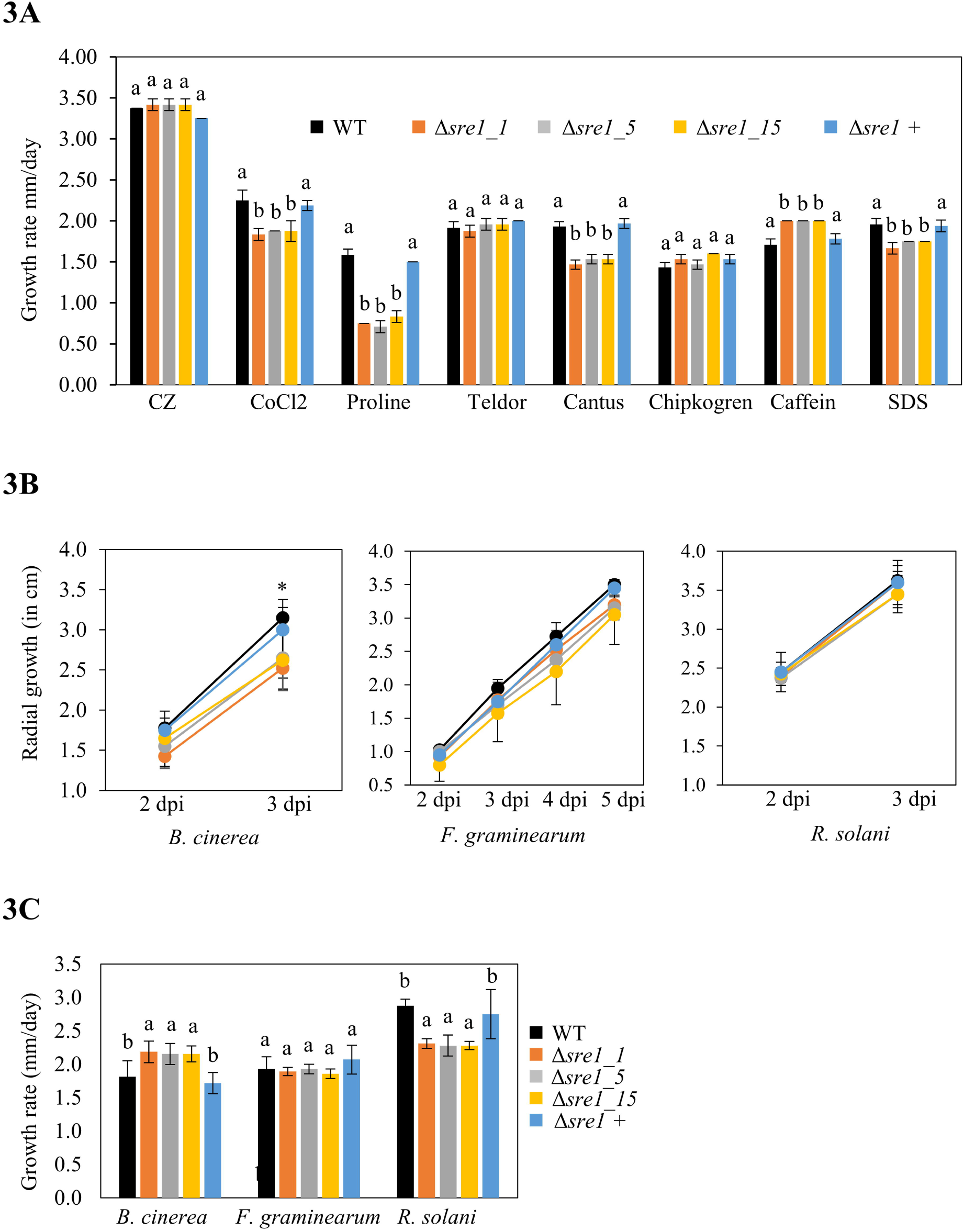
**A**: Growth rate of *C. rosea* WT or Δ*sre1* on czapek-dox medium (CZ) or CZ medium supplemented with fungicides and cell wall stressors. **B**: Colony diameter of mycohosts *B. cinerea*, *F. graminearum*, and *R. solani* during growth in dual culture with *C. rosea* WT and Δ*sre1.* **C**: *C. rosea* WT and Δ*sre1* growth rate over mycohosts in dual culture.

To determine if deletion of *sre* genes affects *C. rosea’*s ability to tolerate hypoxia, the growth rate of the WT and deletion strains was compared on a CZ medium amended with 2.5 mM CoCl_2_. The Δ*sre1* strains displayed a 17% reduced growth rate (*P* < 0.01) on the CoCl2-supplemented CZ medium compared to the WT (**Figure 3A**). Since SREBP regulates ergosterol biosynthesis, we hypothesised that deleting *sre1* and *sre2* would influence *C. rosea* cell wall and membrane integrity. To evaluate this, *C. rosea* strains were grown on CZ supplemented with cell wall stress inducers caffeine or SDS, which has been used as criteria for testing cell wall integrity in fungi and yeasts (Klis et al., 2002; Kuranda et al., 2006; Nunez et al., 2008). The growth rate of Δ*sre1* strains was increased by 17 % (*P* < 0.001) on Caffeine; however, there was a 12 % increase (*P* ≤ 0.007) on the SDS medium (**Figure 3A**). No differences in mycelial growth rate were detected between the WT and Δ*sre2* deletion strains on these media (**Supplementary figure 6**). Furthermore, the complementation strain Δ*sre1*+ showed complete restoration of the growth rate phenotypes observed in Δ*sre1* (**Figure 3A**).

### Deletion of *sre1* altered *C. rosea* antagonism

An in *vitro* dual culture assay was used to test whether deletion of *sre1* or *sre2* affected the antagonistic ability of *C. rosea* against the mycohosts *B. cinerea*, *F. graminearum* and *R. solani*. Three dpi, *B. cinerea* exhibited a significant (*P* ≤ 0.036) 17 % decrease in growth rate during a confrontation with Δ*sre1* strains compared to the WT (**Figure 3B**). In contrast, no differences in the growth rate of *F. graminearum* and *R. solani* were recorded during the same conditions. Similarly, no differences in growth rate between *C*. *rosea* WT and deletion strains were measured during the confrontation with the mycohosts. After the mycelial contact, *C. rosea* strains and the mycohosts were allowed to interact with each other for ten days. The growth rate of Δ*sre1* strains was 20 % higher (*P* ≤ 0.012) on *B*. *cinerea* mycelium (overgrowth rate) compared to the growth rate of WT (**Figure 3D**). In contrast, overgrowth on *R*. *solani* was reduced by 20 % (*P* < 0.001); however, overgrowth on *F. graminearum* was not compromised (**Figure 3C**). Like *in vitro* antagonism tests, a bioassay for biocontrol of fusarium foot rot diseases on wheat caused by *F*. *graminearum* showed no significant difference in biocontrol ability between the WT and the Δ*sre1* strain. Moreover, no differences in antagonism and biocontrol ability were found between the WT and Δ*sre2* (**Supplementary figure 6)**.

### Transcriptome analysis of *C. rosea* WT and *sre1* deletion strain

To dissect the SREBP-mediated gene regulatory network in *C. rosea* and understand the underlying mechanism of impaired Δ*sre1* phenotypes, the transcriptome of *C. rosea* WT and Δ*sre1* was compared in the submerged liquid CZ medium (CZ) and CZ medium supplemented with prothioconazole (CZ + Pro). The sequencing obtained on average 30.9 million reads per sample, 71% of which had unique matches on *C. rosea* genes, while the others went unmapped or were multimapping to several genomic regions or to regions not covered by genes (**Supplementary table 4**). The analysis identified 2145 genes commonly upregulated in Δ*sre1* both treatments compared to the WT, while 544 and 580 genes were uniquely upregulated in CZ and CZ + Pro, respectively. Similarly, 2480 genes were downregulated in both media, while 572 and 704 genes were downregulated in only one of them (**Figure 4A, Supplementary table 5**). The upregulated genes in CZ and CZ + Pro were significantly enriched in 282 and 352 biological processes GO terms, respectively **(Figure 4B; Supplementary table 6**). Among the enriched GO terms, a high proportion was related to metabolic and biosynthetic processes, followed by membrane transport and respiration (**Figure 4C)**. The proportion of enriched biological process GO terms associated with biosynthetic processes increased from 18% in CZ to 21% in the CZ + Pro medium (**Figure 4C**). More specifically, the number of GO terms associated with aa biosynthetic processes, including arginine, glutamine, lysine and *de novo* IMP biosynthetic processes, was higher in CZ + Pro medium compared to CZ. Meanwhile, the number of GO terms associated with aerobic respiration, iron homeostasis, lipid biosynthesis, and pigment biosynthesis was reduced **(Figure 4C**). In contrast to upregulated genes, only one biological process GO term (GO: GO:0006468; protein phosphorylation) was enriched in the genes downregulated in the mutant (**Supplementary table 6**).

**Figure 4:**
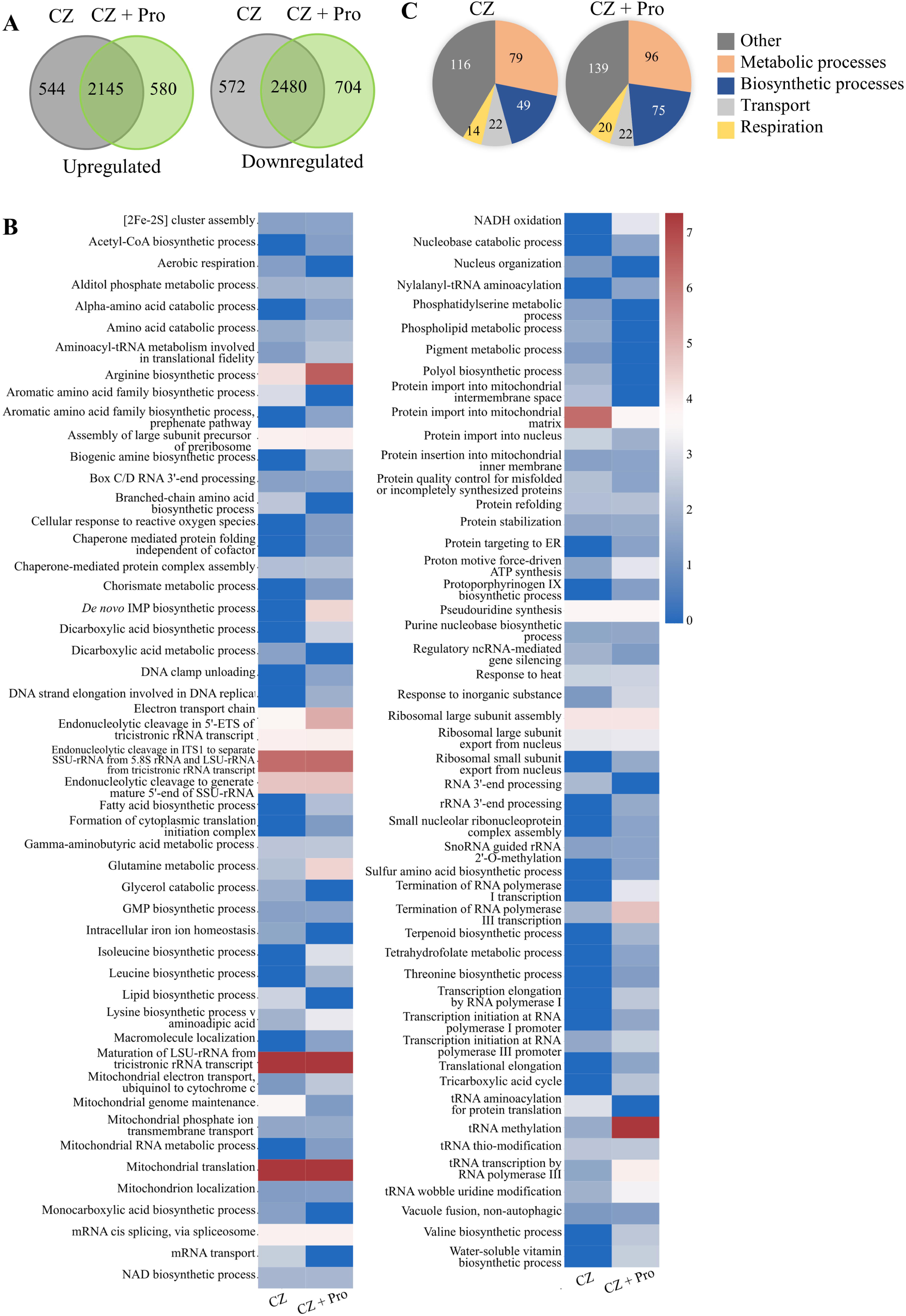
**A:** Number of genes differentially expressed during the transcriptome analysis. **B:** GO terms of biological process enriched in the genes upregulated in Δ*sre1* compared to *C. rosea* WT. When more GO terms described the same processes, only the most specific term was used. The heatmap shows the negative LOG10 of the FDR values determining the significance of the enrichment, calculated with BLAST2GO. **C:** Number of GO terms enriched in the genes upregulated in Δ*sre1* compared to *C. rosea* WT.

To study if deletion of *sre1* affected regulatory feedback loops that influence expression levels of SREBP-associated genes, the expression of *sre2*, *scap* and *insig* was examined by comparing their expression pattern in *C. rosea* WT and Δ*sre1*. The deletion of *sre1* resulted in the downregulation of s*re2,* while the expression of *scap* and *insig* was unaffected (**Table 2**). This suggests that the expression of *sre2* is coregulated with *sre1*, while the expression of *insig* and *scap* is independent of the expression of SREBP in *C. rosea*.

**Table 2:** Expression analysis of *sre2*, *insig* and *scap* genes in the *sre1* deletion strain compared to *C. rosea* WT. Values in bold indicate significant differential expression with FDR < 0.05.

| Gene name | Log2(FC) on CZ | Log2(FC) on CZ + Pro |
| --- | --- | --- |
| *sre2* | **-2.57** | -2.46 |
| *scap* | 0.15 | -0.12 |
| *insig* | 0.92 | 0.67 |

### Deletion of s*re1* triggered transcriptional reprogramming of genes associated with lipid metabolism, aerobic respiration and xenobiotic tolerance

Since SREBPs are shown to regulate gene expression patterns of genes associated with lipid homeostasis, tolerance to hypoxia and drug tolerance, the transcriptome of *C. rosea* strains was further analysed, focusing on genes involved in these processes. Our analysis showed downregulation of monooxygenase coding gene *erg1*, which catalyses the first step in the ergosterol biosynthetic pathway, in Δs*re1* compared to the WT. Intriguingly, seven mevalonate pathway genes (*erg10*, *erg13*, *hmgr*, *erg12*, *idi1*, *erg20*, *erg9*) that biosynthesise precursors of ergosterol and carotenoid, were upregulated (**Figure 5**). Similarly, *crtYB* and *crtI,* part of the carotenoid biosynthetic pathway, and *erg7*, *erg11* and *erg26*, part of ergosterol biosynthetic pathways, were upregulated. In addition, we identified 51 DEGs (29 upregulated and 22 downregulated) in the Δs*re1* compared to the WT with a putative role in lipid biosynthetic and metabolic processes (**Supplementary table 7**).

**Figure 5:**
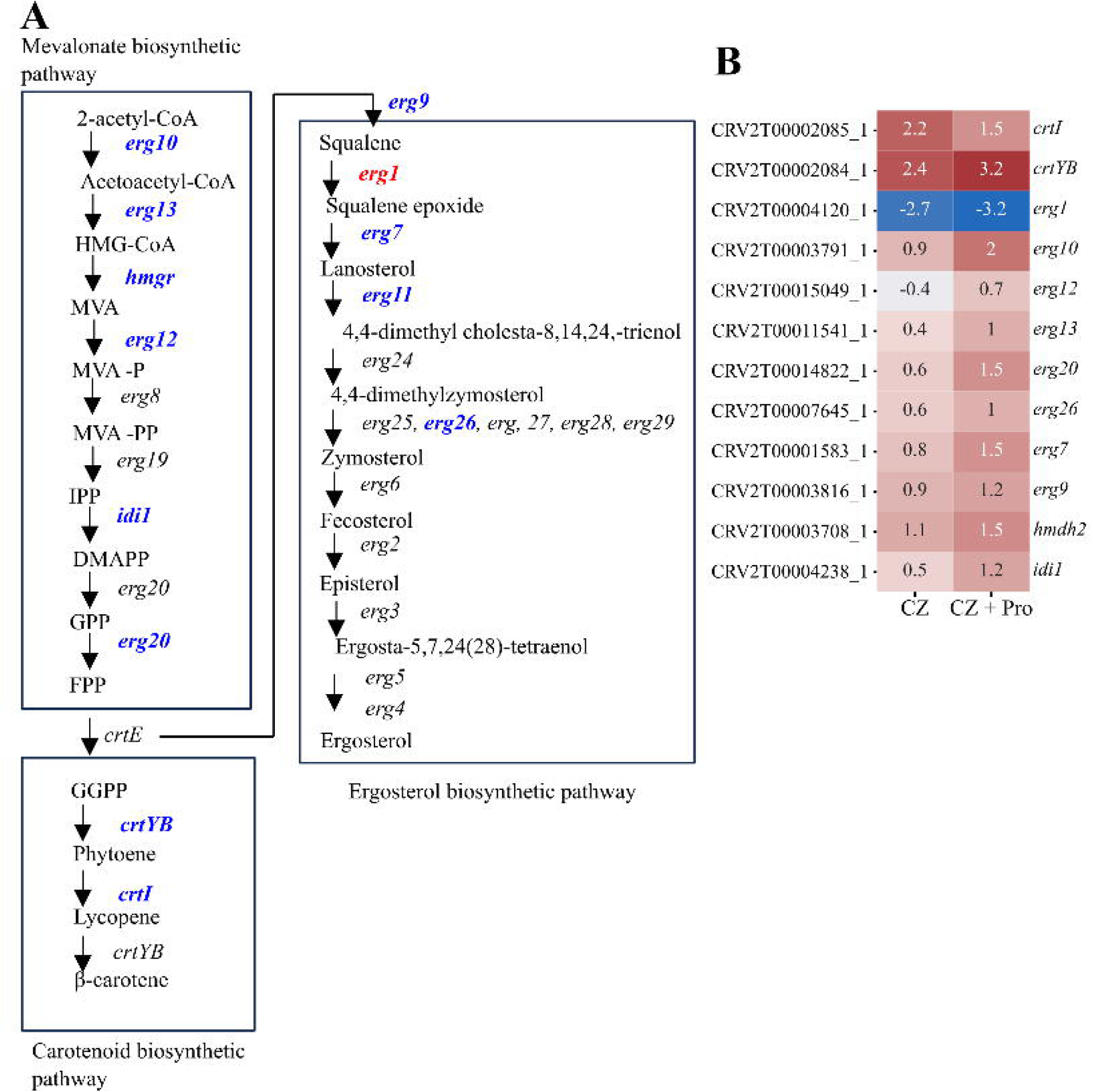
Ergosterol biosynthetic genes differentially expressed in Δ*sre1*. The log2(FC) of the deletion mutant compared to the WT is shown in the heatmap. HMG-CoA, 3-hydroxy-3-methylglutaryl-CoA; MVA, mevalonate; MVA-P, mevalonate-5-phosphate; MVA-PP, mevalonate-5-pyrophosphate; IPP, isopentenyl-pyrophosphate; DMAPP, dimethylallyl-pyrophosphate; GPP, geranyl-pyrophosphate; FPP, farnesyl-pyrophosphate; GGPP, geranylgeranyl-pyrophosphate. crtI, Phytoene desaturase; crtYB, Phytoene synthase; ERG1, Squalene monooxygenase; ERG10, Acetyl-CoA acetyltransferase-like protein; ERG12, Mevalonate kinase; ERG13, Hydroxymethylglutaryl-synthase; ERG20, Farnesyl pyrophosphate synthase; ERG26, Sterol-4-alpha-carboxylate 3-dehydrogenase; ERG7, Terpene cyclase; ERG9, Squalene synthase-like protein; HMDH2, Hmg CoA reductase; IDI1, Isopentenyl-diphosphate delta-isomerase.

We analysed the expression patterns of genes associated with aerobic respiration, such as those involved in glycolysis, tricarboxylic acid (TCA) cycle and mitochondrial electron transport. Seven genes out of ten involved in glycolysis were downregulated in the Δs*re1* compared to the WT (**Figure 6**). Similarly, a gene coding for peroxisomal malate dehydrogenase MDH3, which catalyses the conversion of malate to oxaloacetate in the glyoxylate cycle, an anabolic variant of the TCA cycle, is downregulated in Δs*re1* compared to the WT. The alcohol dehydrogenase (ADH1) gene responsible for catalysing the reduction of acetaldehyde to ethanol during fermentation is also downregulated. In contrast to glycolysis, TCA cycle genes were upregulated in Δ*sre1* except for citrate synthase gene *cit3,* which catalyses the first step of the TCA cycle, performing the condensation of acetyl-CoA with oxaloacetate to form citrate (**Figure 6)**. Similarly, 12 mitochondria genes associated with the electron transport chain and aerobic respiration were upregulated. This includes genes coding for subunits of succinate dehydrogenase (ubiquinone), ubiquinol cytochrome C reductase, cytochrome b-c1 complex, CHCH domain protein and mitochondrial trans-2-enoyl-CoA reductase (**Figure 6**). However, a gene (CRV2T00003883_1) coding for 2-hexaprenyl-6-methoxy-1,4-benzoquinone methyltransferase, required for ubiquinone biosynthesis, was found to be downregulated.

**Figure 6:**
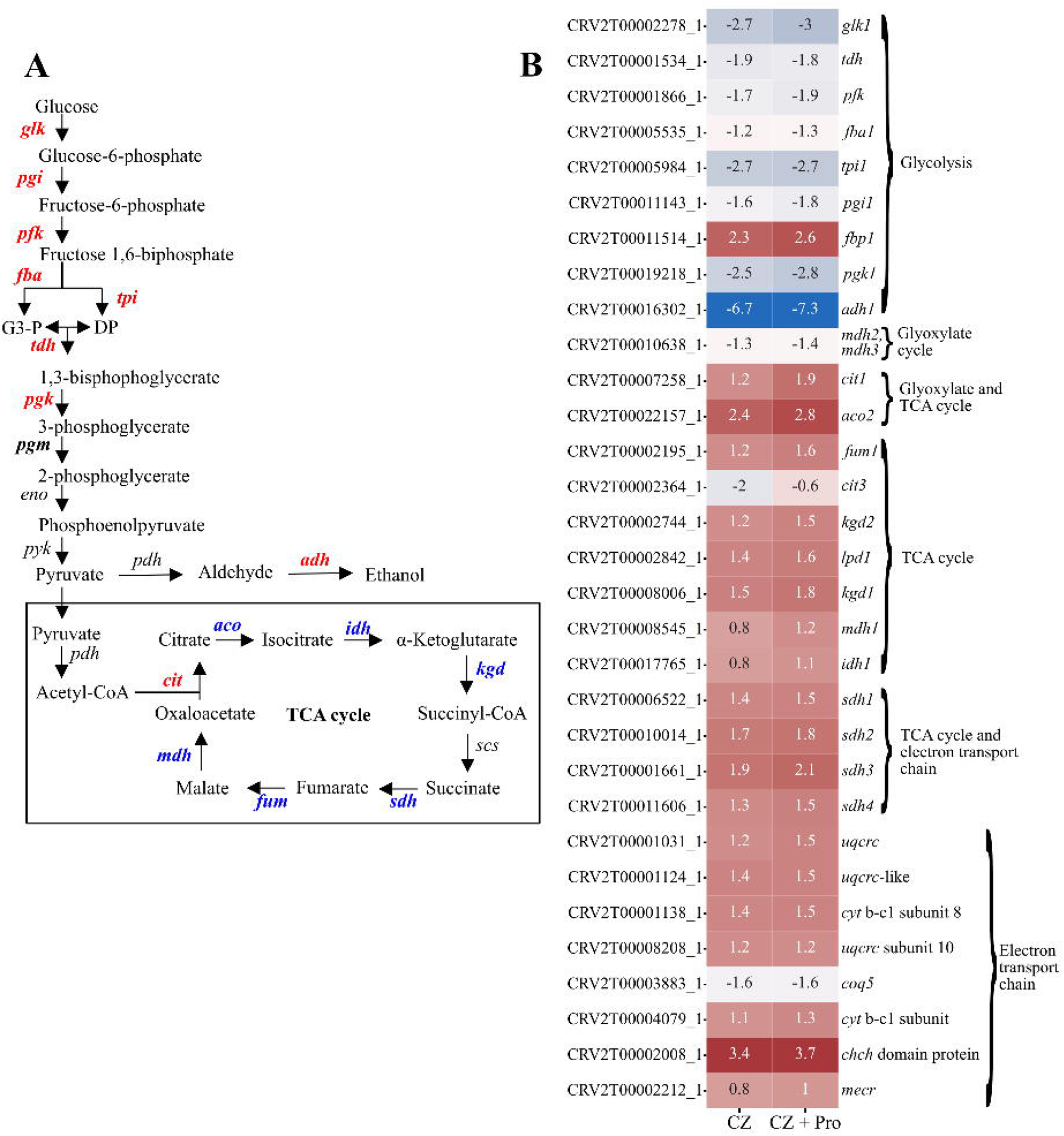
Gene involved in glycolysis, glyoxylate cycle, TCA cycle, or electron transport chain differentially expressed in Δ*sre1*. The log2(FC) of the deletion mutant compared to the WT is shown in the heatmap. GLK, Hexokinase;, PGI, Glucose-6-phosphate isomerase; PFK, Phosphofructokinase; FBA, Aldolase; TPI, Triose phosphate isomerase; TDH, Glyceraldehyde-3-phosphate dehydrogenase; PGK, Phosphoglycerate kinase; PGM, Phosphoglycerate mutase; ENO, Enolase: PYK, Pyruvate kinase; G3-P, Glyceraldehyde 3-phosphate; DP, Dihydroxyacetone phosphate; ADH1, Alcohol Dehydrogenase; ACO, Aconitase; IDH, Isocitrate dehydrogenase; KGD, α-ketoglutarate dehydrogenase; SCS, Succinate-Co-A synthetase; SDH, Succinate dehydrogenase; FUM, Fumarase; MDH, malate dehydrogenase; CIT, Citrate synthase; PDH, Pyruvate dehydrogenase; PDC, Pyruvate decarboxylase.

Our analysis showed 38 DEGs (17 upregulated, 21 downregulated) related to iron homeostasis, which plays an essential role in cellular respiration and oxygen transport in fungi (Blatzer et al., 2011). This includes 13 genes coding for heme peroxidase, iron permease FTR1 family protein, siderophore iron transporter mirb protein, FeS biogenesis, and iron-sulfur cluster assembly protein (**Figure 7**). We also identified 13 differentially expressed MFS transporter genes with high similarity with siderophore-iron transporter 1 *Sit1* and *Str1*, ferri-siderophore transporter *MirB* and Fusarium iron-related protein *Fir1* (**Figure 7**).

**Figure 7:**
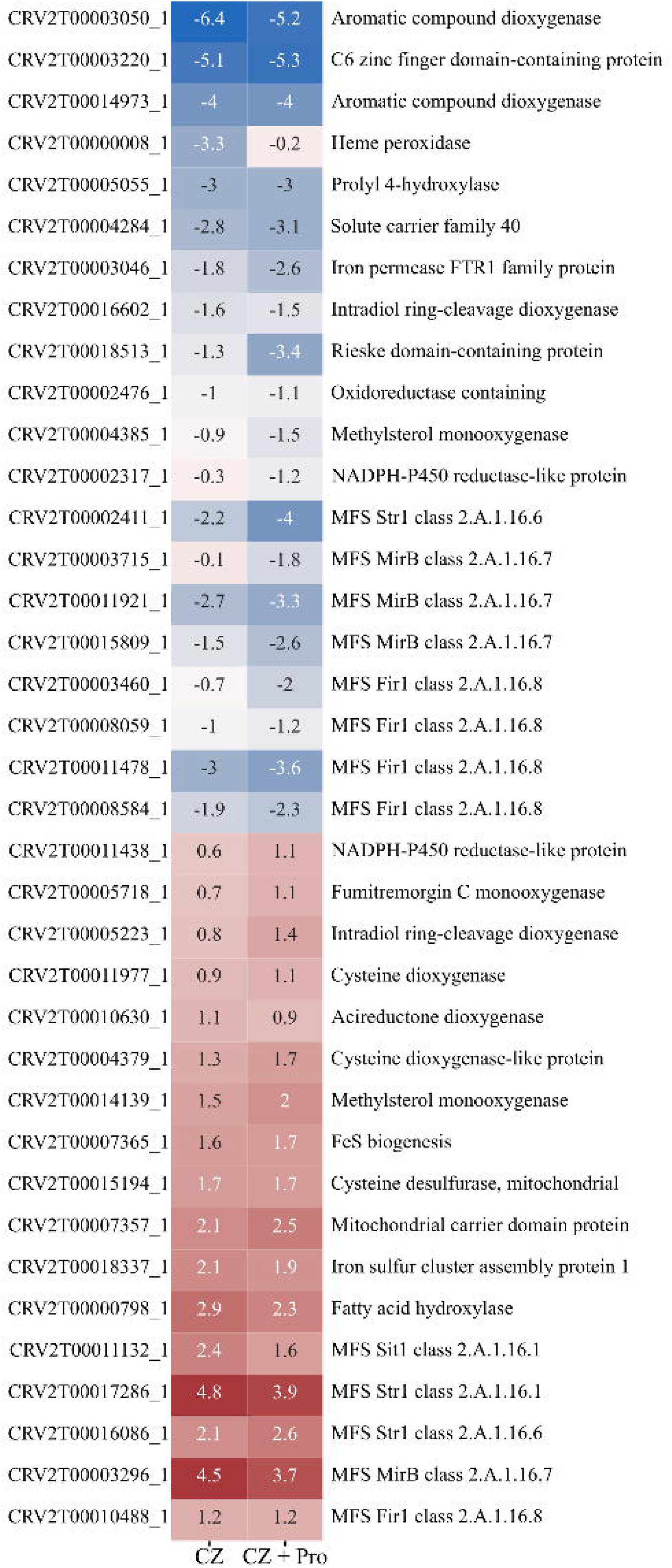
Genes involved in iron homeostasis differentially expressed in Δ*sre1*. The log2(FC) of the deletion mutant compared to the WT is shown in the heatmap. HS, *Helminthosporium solanum*; FG, *Fusarium graminearum*; BC, *Botrytis cinerea*.

To test the role of membrane transporters in fungicide tolerance in *C. rosea*, we compared the expression pattern of membrane transporter genes between Δ*sre1* and the WT strains. We focused our analysis on MFS (major facilitator superfamily) transporters and ABC (ATP-binding cassette) transporter genes, which are considered crucial for their role in fungicide tolerance in *C. rosea* (Broberg et al., 2018; Dubey et al., 2014a, 2016; Funck Jensen et al., 2021; Karlsson et al., 2015). Gene expression analysis identified 215 MFS transporter genes differentially expressed in Δ*sre1* compared to the WT. Among these, 52 (34 upregulated 18 downregulated) were classified as putative drug transporters with potential roles in fungicide and xenobiotic tolerance (**Table 3, supplementary table 7**). This includes genes showing high similarity with MFS transporters *mfs1, qdr2* and *qdr3*, which have been characterised for their role in drug, mycotoxins and fungicide tolerance (Liu et al., 2012; Qadri et al., 2022; Saier Jr et al., 2021). Similarly, 49 ABC transporter (30 upregulated and 19 downregulated) genes were differentially expressed. This include 15 from the pleiotropic drug resistance protein (PDR) family ABC-G, seven from multidrug resistance-associated protein (MRP) family ABC-C and ten from multidrug resistance protein (MDR) family ABC-B were differentially expressed in Δ*sre1* compared to the WT (**Table 3, Supplementary table 7**). The expression pattern of genes coding for Cytochrome p450 (CYP), a heme-containing protein involved in xenobiotic metabolism in fungi, was also analysed (Chen et al., 2014; Manikandan and Nagini, 2018). Our analysis identified 75 differentially expressed (31 upregulated and 44 downregulated) CYP450 genes in Δ*sre1* compared to the WT (**Supplementary table 7**). These include CYP51 gene coding for eburicol 14-alpha-demethylase-like protein and isotrichodermin C-15 hydroxylase, critical in the ergosterol biosynthesis pathway and secondary metabolite production.

**Table 3:** number and class of MFS and ABC transporters differentially expressed in Δ*sre1*. MFS transporters were classified according to their TCDB role (Saier Jr et al., 2021), while MFS transporters were classified according to the classification of Kovalchuk and Driessen (2010). CZ: czapek-dox medium; CZ+Pro: czapek-dox medium with the addition of prothioconazole.

| MFS class | TCDB Role | Upregulated<br>in CZ | Downregulated<br>in CZ | Upregulated<br>in CZ+Pro | Downregulated<br>in CZ+Pro |
| --- | --- | --- | --- | --- | --- |
| 2.A.1.1 | Sugar Porters<br>(SP) | 18 | 24 | 13 | 27 |
| 2.A.1.12 | Sialate:H+<br>Symporters (SHS) | 0 | 0 | 0 | 1 |
| 2.A.1.13 | Monocarboxylate<br>Transporters<br>(MCT) | 10 | 7 | 9 | 6 |
| 2.A.1.14 | Anion:Cation<br>Symporters<br>(ACS) | 12 | 15 | 13 | 11 |
| 2.A.1.16 | Siderophore-Iron<br>Transporters<br>(SIT) | 5 | 5 | 4 | 5 |
| 2.A.1.19 | Organic Cation<br>Transporters<br>(OCT) | 1 | 2 | 2 | 3 |
| 2.A.1.2 | Drug:H+<br>Antiporters-1 (12<br>Spanner) (DHA1) | 28 | 17 | 28 | 15 |
| 2.A.1.3 | Drug:H+<br>Antiporters-2 (14<br>Spanner) (DHA2) | 14 | 12 | 13 | 10 |
| 2.A.1.48 | Vacuolar Basic<br>Amino Acid<br>Transporters (V-BAAT) | 2 | 0 | 3 | 0 |
| 2.A.1.53 | Proteobacterial<br>Intraphagosomal<br>Amino Acid<br>Transporters (Pht) | 0 | 0 | 1 | 0 |
| 2.A.1.7 | Fucose: H+<br>Symporters (FHS) | 0 | 1 | 0 | 3 |
| 2.A.1.9 | Phosphate: H+<br>Symporters (PHS) | 1 | 1 | 1 | 2 |
| ABC class | Description of the<br>class<br>(Kovalchuk and<br>Driessen (2010)) |  |  |  |  |
| FG_ABC_A | Lipid transport<br>and metabolism | 0 | 1 | 0 | 1 |
| FG_ABC_B | Multidrug<br>resistance | 9 | 5 | 6 | 5 |
| FG_ABC_C | Detoxification of<br>toxic compounds | 4 | 4 | 4 | 3 |
| FG_ABC_F | Translation, not<br>transport | 2 | 0 | 4 | 0 |
| FG_ABC_G | Pleiotropic drug | 9 | 6 | 8 | 4 |

|  | resistance |  |  |  |  |
| --- | --- | --- | --- | --- | --- |
| FG_ABC_ICaf16p | Unknown | 2 | 0 | 2 | 0 |
| FG_ABC_Ydr061r | Unknown | 1 | 0 | 1 | 0 |

### Antagonism-responsive genes were downregulated in **Δ***sre1* strain

To investigate if the altered antagonistic ability of Δ*sre1* is due to a change in the expression of antagonism-responsive *C. rosea* genes, we used transcriptome data from other works collected during *C. rosea* interactions with plant pathogenic mycohosts *B. cinerea*, *F. graminearum* and *Helminthosporium solani* (Demissie et al., 2020, 2018; Lysøe et al., 2017; Nygren et al., 2018; Piombo et al., 2021). *C. rosea* genes upregulated during interactions with mycohosts were considered antagonism-related genes in this study. In total, we found 27 differentially expressed antagonism-related genes in the Δ*sre1*, compared to the WT. Of these, 18 genes were downregulated in Δ*sre1* compared to the WT, including genes coding for polyketide synthases (*pks22*, *pks23*, *pks29*), MFS transporters (*mfs104*, *mfs524*, *mfs533*), proteases, peptidases, chitinases and Cathepsin A (**Figure 8**).

**Figure 8:**
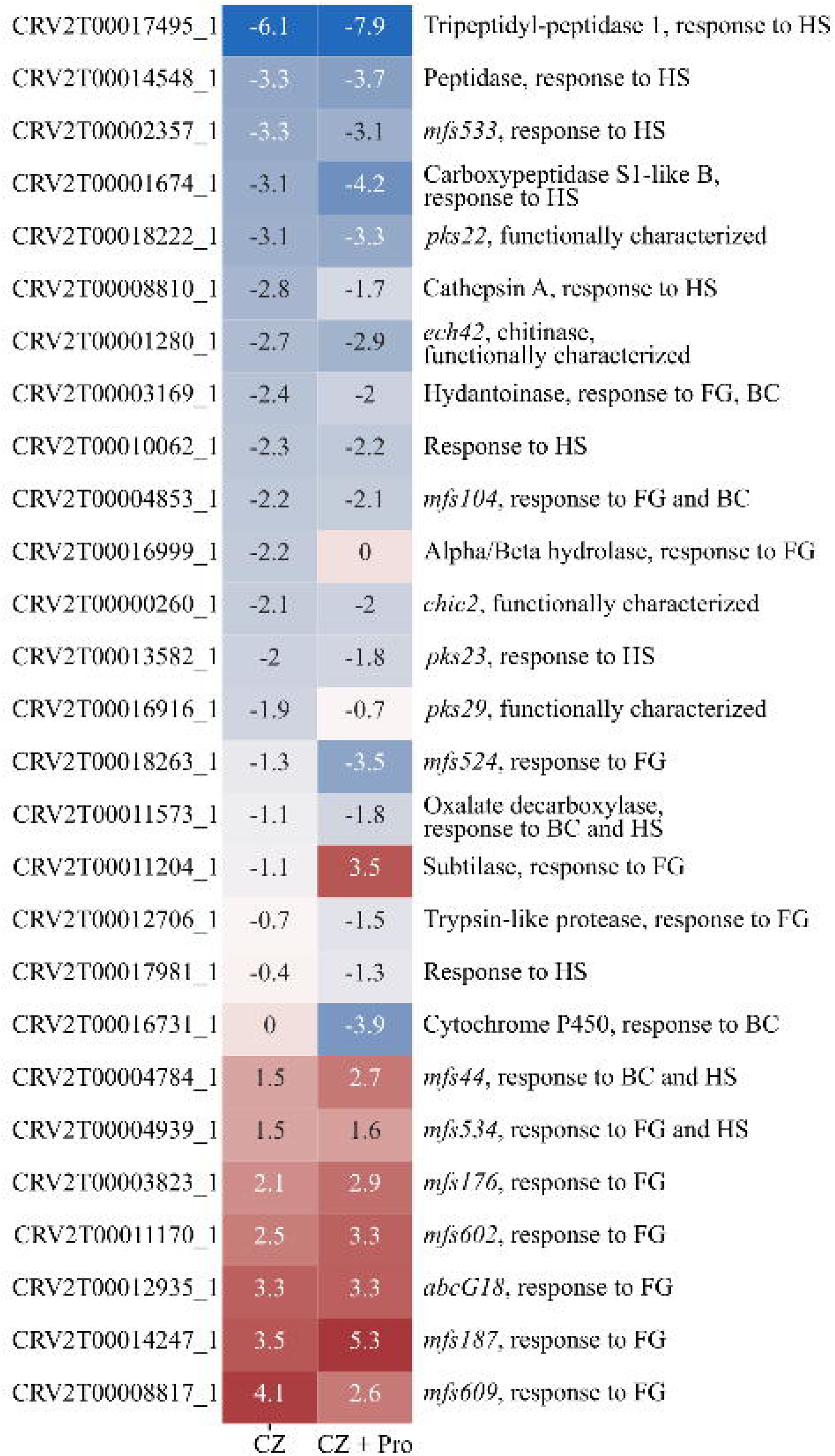
Genes involved in response to plant pathogens differentially expressed in Δ*sre1*. The log2(FC) of the deletion mutant compared to the WT is shown in the heatmap. HS, *Helminthosporium solanum*; FG, *Fusarium graminearum*; BC, *Botrytis cinerea*.

## Discussion

SREBPs have been functionally characterized for their roles in sterol biosynthesis in mammals (Osborne and Espenshade, 2009). The SREBP orthologs in the fungal kingdom have been identified and described in fungal species with diverse lifestyles, including yeast *S. pombe*, human plant pathogenic fungi such as *Cryptococcus neoformans* and *A. fumigatus*, and plant pathogenic fungi such as *M. oryzae* and *Penicillium digitatum* (Chung et al., 2019; Osborne and Espenshade, 2009; Ruan et al., 2017). Here, we provided significant insights into the biological and regulatory roles of SREBPs in the mycoparasitic fungus *C. rosea* with a focus on biocontrol traits, fungicide tolerance and adaptation to hypoxia.

Genes coding for SREBPs are variably distributed among fungal kingdoms and are broadly divided into SRE clade A (SreA or clade 1) and clade B (SreB or clade 2) (Ruan et al., 2019). Among fungal species belonging to class Sordariomycetes in Ascomycota, for instance, plant pathogens and saprotrophs contain three genes, while animal pathogens possess only one gene in their genome coding for SREBP (Chung et al., 2019; Ruan et al., 2019). However, in Basidiomycota, the distribution of SREBP genes is patchy, with no SREBP genes in many fungal species, including plant-pathogens *Puccinia graminis* and *Melampsora laricis-populina*, human pathogen *C. neoformans*, root endophyte *Piriformospora indica* and mycorrhizae *Laccaria bicolor,* while one gene is present in saprotroph *Phanerochaete chrysosporium* and mycorrhizae *Piloderma croceum* (Chung et al., 2019). This SREBP gene copy numbers diversity is plausibly related to fungal adaptations to different ecological niches, and for this reason we analysed the distribution of gene copy numbers, focussing on the fungal species in Hypocreales. The results showed a similarity in the gene copy number distribution of SREBP in the clade SreA among fungal species with diverse lifestyles, indicating a conserved regulatory mechanism of ergosterol biosynthesis across Hypocreales (Ruan et al., 2019). We identified two SREBPs, SRE1 and SRE2, in *C. rosea* with typical SREBP characteristics, belonging to SRE clade one and SRE clade two, respectively (Osborne and Espenshade, 2009; Ruan et al., 2019). The lack of C terminus DUF2014 domain is a characteristic of SRE clade 2, and the domain was missing in *C. rosea* SRE2, similarly to previously characterised SREBPs such as MoSRE2 and MoSRE3 of *M. oryzae*, and SRBB of *A. fumigatus* (Chung et al., 2019; Gutiérrez et al., 2019; Maguire et al., 2014). Another characteristic of SREBPs is a transmembrane helix that anchors them to the ER membrane (Osborne and Espenshade, 2009). Like *A. fumigatus* SRBA, *M. oryzae* MoSRE1 and *S. pombe* SRE1 and SRE2 (Chung et al., 2019; Osborne and Espenshade, 2009), *C. rosea* SRE1 contain two transmembrane helices. However, the lack of transmembrane helix in *C. rosea* SRE2 is similar to *A. fumigatus* SRBB, and other SREBPs encoded by *N. crassa*, *C. neoformans* and *M. oryzae* (Chung et al., 2019; Maguire et al., 2014; Qin et al., 2017). Phylogenetic distribution and protein structure differences between SRE1 and SRE2 indicate a functional diversification of SREBP in *C. rosea*. In addition to SREBPs, identifying INSIG and SCAP coding genes, which are essential components of SREBP-mediated sterol homeostasis, indicates the existence of the SREBP signaling pathway in *C. rosea* (Osborne and Espenshade, 2009). This is supported by our results, showing the physical interaction between *C. rosea* SRE1 and SCAP. No physical interaction between SCAP and SRE2 suggests that the activation mechanism of *C. rosea* SRE1 differs from that of SRE2 and corroborates the role of the DUF2014 domain in SREBP-SCAP interaction. Taken together, our data showed the presence of the genetic machinery required for the SREBP signaling pathway in *C. rosea* and highlighted that SRE1 and SRE2 belong to phylogenetically and structurally divergent groups of SREBP and have evolved for different roles in *C. rosea*.

The SREBP-mediated regulatory mechanisms of sterol biosynthesis are primarily conserved between animals and Ascomycetes fungi (Bien and Espenshade, 2010; Osborne and Espenshade, 2009). However, certain fungal species, including *S. cerevisiae* and *C. albicans,* lack mammalian SREBP homologues and have evolved distinct regulatory mechanisms. In *S. cerevisiae,* for instance, the sterol biosynthesis pathway is regulated by another regulatory protein, Upc2, which is a Gal4-type zinc finger transcription factor (Butler, 2013; Maguire et al., 2014; Ruan et al., 2019). Although Upc2 and SREBPs are structurally different, they are localized to intracellular membranes and share functional similarities (Butler, 2013), including tolerance to hypoxia and azoles (Liu et al., 2015; Osborne and Espenshade, 2009). In the current study, heterologous expression of *C. rosea sre1* in Δ*Upc2* strain background failed to rescue *S. cerevisiae* from prothioconazole toxicity, which is in line with previous findings suggesting SREBP-mediated sterol regulatory mechanism is unrelated to that regulated by Upc2 and demonstrating evolutionary diversity in sterol regulatory mechanisms among fungi (Liu et al., 2015; Osborne and Espenshade, 2009). However, enhanced tolerance of *S. cerevisiae* WT expressing *C. rosea sre1* to prothioconazole highlights the potential role of SRE1 in conferring tolerance to this fungicide in *C. rosea*. This result also underscores the functional diversification between SRE1 and SRE2 in *C. rosea*, as *sre2* overexpression did not affect *S. cerevisiae’s* tolerance to prothioconazole. This is corroborated by the results of gene deletion, where no significant differences in the tested phenotypes were observed between the WT and Δ*sre2*, while Δ*sre1* showed several phenotypic effects.

In line with previous findings (Blatzer et al., 2011; Chung et al., 2019; Liu et al., 2015; Osborne and Espenshade, 2009; Ruan et al., 2017; Willger et al., 2008), deletion of *C. rosea sre1* resulted in phenotypic effects including lower growth rates on medium supplemented with chemical compounds, targeting ergosterol biosynthesis and fungal respiration. This phenotypic effect plausibly results from the impaired cell wall function in Δ*sre1* strains under stress conditions. This is justified by the altered growth rate of Δ*sre1* strains on medium supplemented with cell wall stressors and deformation of the mycelial structure on medium supplemented with proline. Since SREBP is a positive regulator of ergosterol biosynthesis genes, the upregulation of many of these genes in SRE1 deletion strains is intriguing and plausibly suggests an additional mechanism of gene expression regulation in response to prothioconazole in *C. rosea*. This also indicates that deletion of *sre1* forms a negative transcriptional feedback loop, resulting in the upregulation of ergosterol biosynthesis genes to maintain cellular homeostasis.

Membrane transporters such as ABC and MFS can mediate the efflux of a wide range of drugs and fungicides across the membrane (Coleman and Mylonakis, 2009; Lamping et al., 2010), and upregulation of this class of membrane transporters is one of the nontarget-site mechanisms of drug and fungicides including azole resistance in fungi (Hu and Chen, 2021; Yin et al., 2023). Similarly, CYP, a membrane-bound heme protein, catalyses many reactions involved in drug metabolism, lipids biosynthesis, and iron homeostasis, thereby playing a pivotal role in fungal secondary metabolisms and xenobiotic/drug detoxification (Chen et al., 2014; Manikandan and Nagini, 2018). Results from comparative transcriptome analysis showing downregulation of many of ABC transporters, MFS transporters and CYP genes in Δ*sre1* strains underpin the role of this group of proteins in tolerance of prothioconazole and fenhexamid in *C. rosea*. Among the downregulated MFS transporters, we identified 11 genes showing similarity with MFS1 previously characterized for its role in fungicide tolerance in the plant pathogenic fungi *Botrytis cinerea*, *Mycosphaerella graminicola*, human pathogenic fungus *Trichophyton rubrum* and the fungal biocontrol agent *Trichoderma harzianum* (Liu et al., 2012; Roohparvar et al., 2007; Samaras et al., 2021; Yamada et al., 2021). We identified three downregulated ABC transporters genes (*abcB4* and *abcB18* from group B, multidrug-resistant; abcC14 from group C, multidrug resistance-associated proteins) in Δ*sre1,* responsive to culture filtrates from biocontrol bacteria *Pseudomonas chororaphis* and *Serratia rubidaea* S55 (Kamou et al., 2016; Karlsson et al., 2015), indicating a potential role in xenobiotic tolerance as culture filtrates often contain secondary metabolites. CYPs are another gene family often involved in xenobiotic tolerance. In *T. atroviride*, the expression of this gene family was induced in the presence of pesticide dichlorvos, and functional characterisation of Ta*Cyp*548-2 by gene deletion demonstrated its involvement in dichlorvos degradation (Hayashi et al., 2002; Nakaune et al., 1998; Omrane et al., 2015; Sun et al., 2022; Whaley et al., 2018; Zhang et al., 2012). The role of these gene families in *C. rosea* fungicide resistance is also supported by the fact that the *C. rosea* genome contains a significantly higher number of genes coding for ABC transporters, MFS transporters and CYPs, considered to be involved in the efflux and metabolism of xenobiotics (Dubey et al., 2016, 2014a; Kamou et al., 2016; Karlsson et al., 2015; Nygren et al., 2018). Deletion of pleiotropic drug resistance transporter ABCG5 and ABCG29 genes, for example, resulted in *C. rosea* strains with reduced ability to tolerate xenobiotics, including fungicides (Dubey et al., 2014a, 2016). Together, these results highlight that the nontarget-site mechanism is one of the crucial traits of intrinsic fungicide resistance in *C. rosea*.

In addition to drug tolerance, the fungal SREBPs are shown to be involved in hypoxia adaptation, iron, heme and lipid homeostasis (Osborne and Espenshade, 2009; Liu et al., 2015; Willger et al., 2008; Chung et al., 2014). Sterol biosynthesis is an oxygen-dependent process that requires a relatively higher amount of oxygen, and several oxygen-dependent enzymes are iron-containing; therefore, responses to lipid and carbohydrate metabolism, hypoxia and iron homeostasis are often coregulated (Chung et al., 2014; Liu et al., 2015; Osborne and Espenshade, 2009; Willger et al., 2008). Reduced growth of Δ*sre1* in CoCl_2_ supplemented medium aligns with these previous findings. Downregulation of genes involved in glycolysis and fermentation and upregulation of genes involved in the TCA cycle and mitochondrial electron transport chain showed SRE1 as a positive regulator of glycolysis in *C. rosea*. It is possible that the upregulation of the TCA cycle in the mutant is due to a negative feedback regulation mechanism caused by the reduced abundance of glycolysis-generated pyruvate. The TCA cycle is strongly connected to both lipid and amino acid metabolism, as lipid synthesis requires citrate, a TCA cycle intermediate, to generate glycerol 3-phosphate and acetyl-CoA (Chandel, 2021a), while another TCA cycle intermediate, α-ketoglutarate, is fundamental to synthetize multiple amino acids (Chandel, 2021b). The upregulation of the TCA cycle genes in *sre1* deletion strain could be the result of reprogramming of the respiratory metabolic pathway for energy production and redox homeostasis. This explanation is supported by the fact that the upregulated genes were significantly enriched in GO terms associated with glycerol catabolic, lipids biosynthetic processes and phospholipids metabolic processes. Similarly, various GO terms related to amino acid biosynthetic and catabolic processes, including glutamine, leucine, isoleucine, and lysine, were also enriched. Our result is in line with previous findings in *A. fumigatus*, where comparative gene expression analysis between WT and SREBP deletion strains showed the role of SREBP in regulating carbohydrate metabolism (Chung et al., 2014). However, previous studies on *S. pombe* and *C. neoformans* showed that SREBPs are a positive regulator of gene expression of anaerobic pathway genes and are mainly redundant for genes involved in glycolysis and TCA cycle (Bien and Espenshade, 2010; Chang et al., 2007; Chun et al., 2007; Todd et al., 2006).

Studies in *A. fumigatus* and *C. neoformans* have shown that SREBPs are required for the adaptation to iron homeostasis (Blatzer et al., 2011; Chang et al., 2007). In line with the previous studies, this work showed downregulation of many genes associated siderophore transport and heme biosynthesis in Δ*sre1*, validating the role of SREBP in iron homeostasis in fungi. Among the downregulated genes, for example, we found *C. rosea* genes with high similarity with siderophore transporters *mirB*, *fir1* and *sit1*, characterised for their role in iron uptake, as well as the iron permease *ftri*, coding for an iron-sulfur protein (Rieske domain protein) which is one of the catalytic subunits of the cytochrome *bc*_1_ complex (Conte and Zara, 2011; Heymann et al., 2002; Park et al., 2006; Raymond-Bouchard et al., 2012; Stearman et al., 1996). Similarly, altered expression of genes associated with oxygen-requiring lipid metabolic processes further validates the role of *C. rosea* SRE1 as a gene expression regulator required for lipid homeostasis.

Production and tolerance of secondary metabolites is vital for *C. rosea*’s antagonistic ability (Broberg et al., 2021; Dubey et al., 2014a; Fatema et al., 2018; Karlsson et al., 2015; Nygren et al., 2018). Our analysis showed down-regulation of mycohost-responsive polyketide synthase (PKS) genes *(pks22*, *pks23*, *pks29*) together with several MFS transporters (*mfs533*, *mfs104*, and *mfs524*) in Δ*sre1*. The *pks22* and *pks29* genes were shown to be highly induced during antagonistic *C. rosea*’s interaction with *B. cinerea* and *F. graminearum*, and deletion of the *pks29* resulted in mutants with a reduced biocontrol ability to control fusarium foot rot on barley (Fatema et al., 2018). Moreover, PKS22 is shown to be required to produce the polyketides clonoroseins A-D, with an antagonistic ability against *B. cinerea* and *F. graminearum* (Fatema et al., 2018). In addition, we detected in Δ*sre1* the downregulation of two chitinase genes, *chic2* and *ech42*, which were previously characterised for their antagonistic ability against *B. cinerea*, *R. solani* and *F. graminearum*, respectively (Mamarabadi et al., 2008; Tzelepis et al., 2015). These results suggest that the downregulation of these genes in Δ*sre1* plausibly influenced the *C. rosea* antagonisms against *R. solani*. The enhanced antagonistic ability of *sre1* deletion strains against *B. cinerea* could instead be explained by the fact that many ABC and MFS transporters were upregulated in the Δ*sre1*, and many genes of this class have been observed to be important in the antagonistic activity of *C. rosea* (Demissie et al., 2020; Nygren et al., 2018).

## Conclusion

We characterized the biological and regulatory role of SREBPs in the fungal BCA *C. rosea*. We identified two SREBP encoding genes *sre1* and *sre2* in *C. rosea* and showed that they are phylogenetically and structurally different and have evolved for different roles. The evidence for the functional diversification between SRE1 and SRE2 is provided by the results of the overexpression in *S. cerevisiae* and the gene deletions, where Δ*sre1* showed several pleiotropic effects, while no significant differences in the tested phenotypes were observed between the WT and Δ*sre2*. The reduced growth rate of *sre1* deletion mutants on medium supplemented with the hypoxia mimicking agent CoCl_2_, or the fungicides proline and Cantus, corroborate the role of SREBPs in intrinsic fungicide tolerance and hypoxia tolerance. In addition, we showed that SRE1 has a role in antagonism towards *B. cinerea* and *R. solani*. By comparing *C. rosea* WT and Δ*sre1* transcriptome, we provided insights into the regulatory role of this gene. Deletion of *sre1* caused reprogramming of gene expression patterns of genes associated with carbohydrate and lipid metabolism, respiration, iron homeostasis, and xenobiotic tolerance. Downregulation of antagonism-responsive genes such polyketide synthase and chitinase genes provided valuable support to the impaired antagonism phenotype of Δ*sre1* strains against *R. solani*. Taken together, the result presented in this study provide valuable insights into the diverse roles of SREBPs in fungal BCAs, offering a deeper understanding of their contributions to biocontrol traits, and adaptive responses to environmental stresses including hypoxia and fungicide tolerance. Exploring the underlying mechanisms of intrinsic fungicide tolerance and biocontrol traits in BCAs is important to combine the application of fungicides and BCAs, favouring the knowledge-based implementation of IPM strategies in agriculture production systems.

## Data availability statement

This paper’s sequencing data is available on ENA under the bioproject PRJEB61889.

## Acknowledgement

This work was financially supported by the Department of Forest Mycology and Plant Pathology. MD acknowledge Swedish Research Council for Environment, Agricultural Sciences and Spatial Planning (FORMAS; grant number 2018-01420, 2021-01461), and The Swedish research council, Vetenskapsrådet (VR; grant number 2022-03639). Sequencing was performed by the SNP&SEQ Technology Platform in Uppsala. The facility is part of the National Genomics Infrastructure (NGI) Sweden and Science for Life Laboratory. The SNP&SEQ Platform is also supported by the Swedish Research Council and the Knut and Alice Wallenberg Foundation.

