## Supplementary figures and images for "SREBP-mediated gene expression regulation is essential for the intrinsic fungicide tolerance and antagonism in the fungal biocontrol agent *Clonostachys rosea*"

### Supplementary figure 1

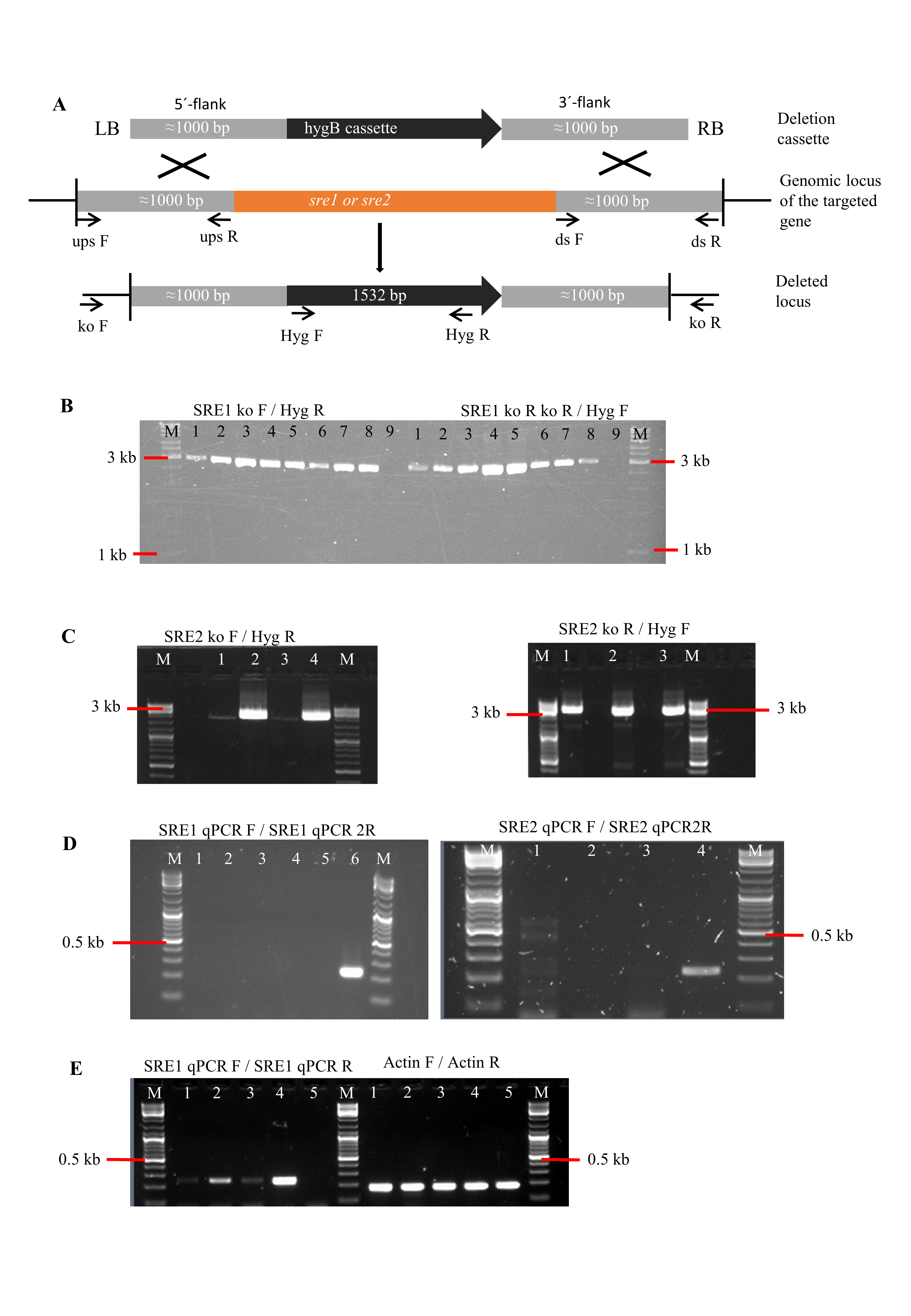

### Supplementary figure 2

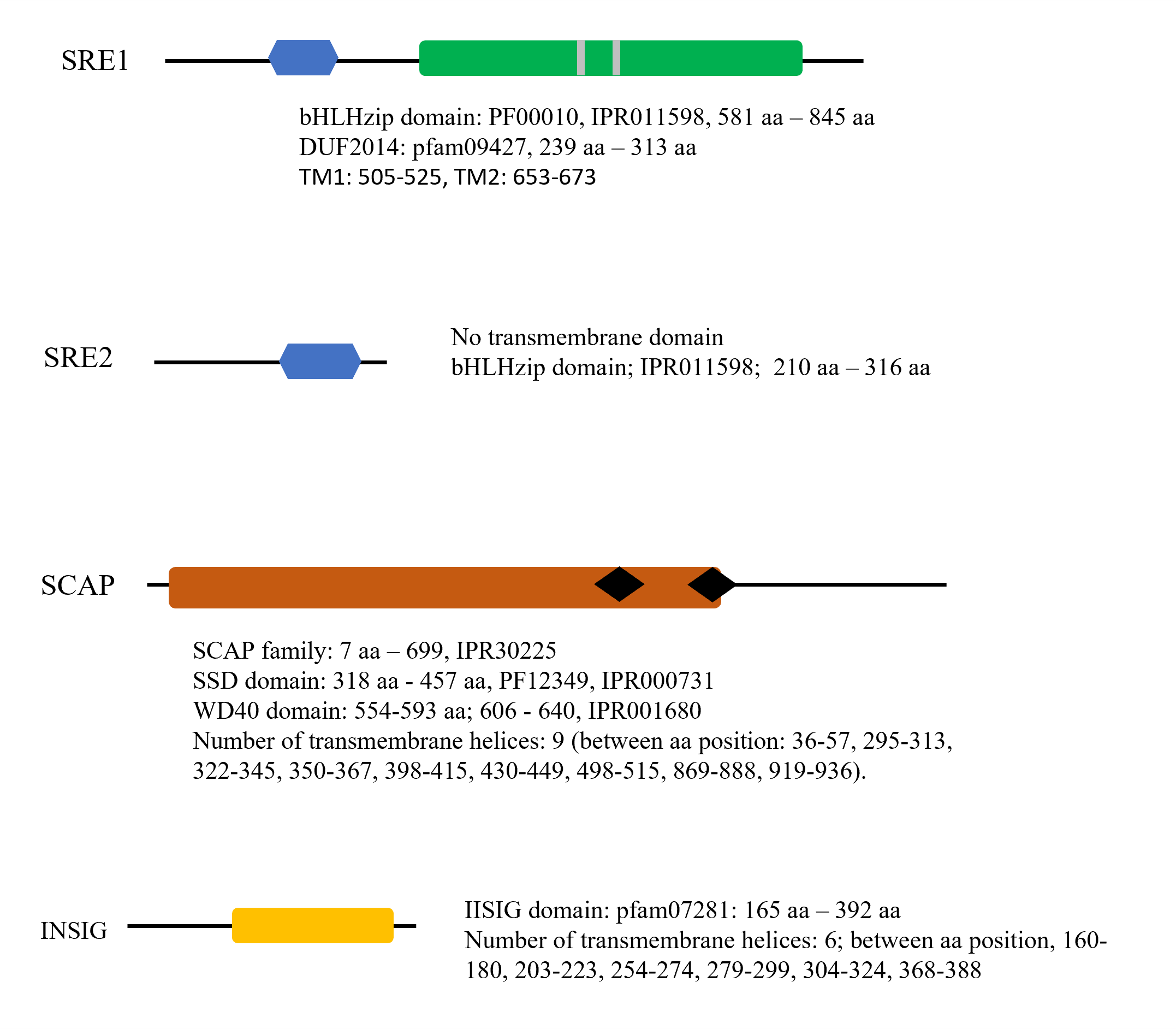

### Supplementary figure 3

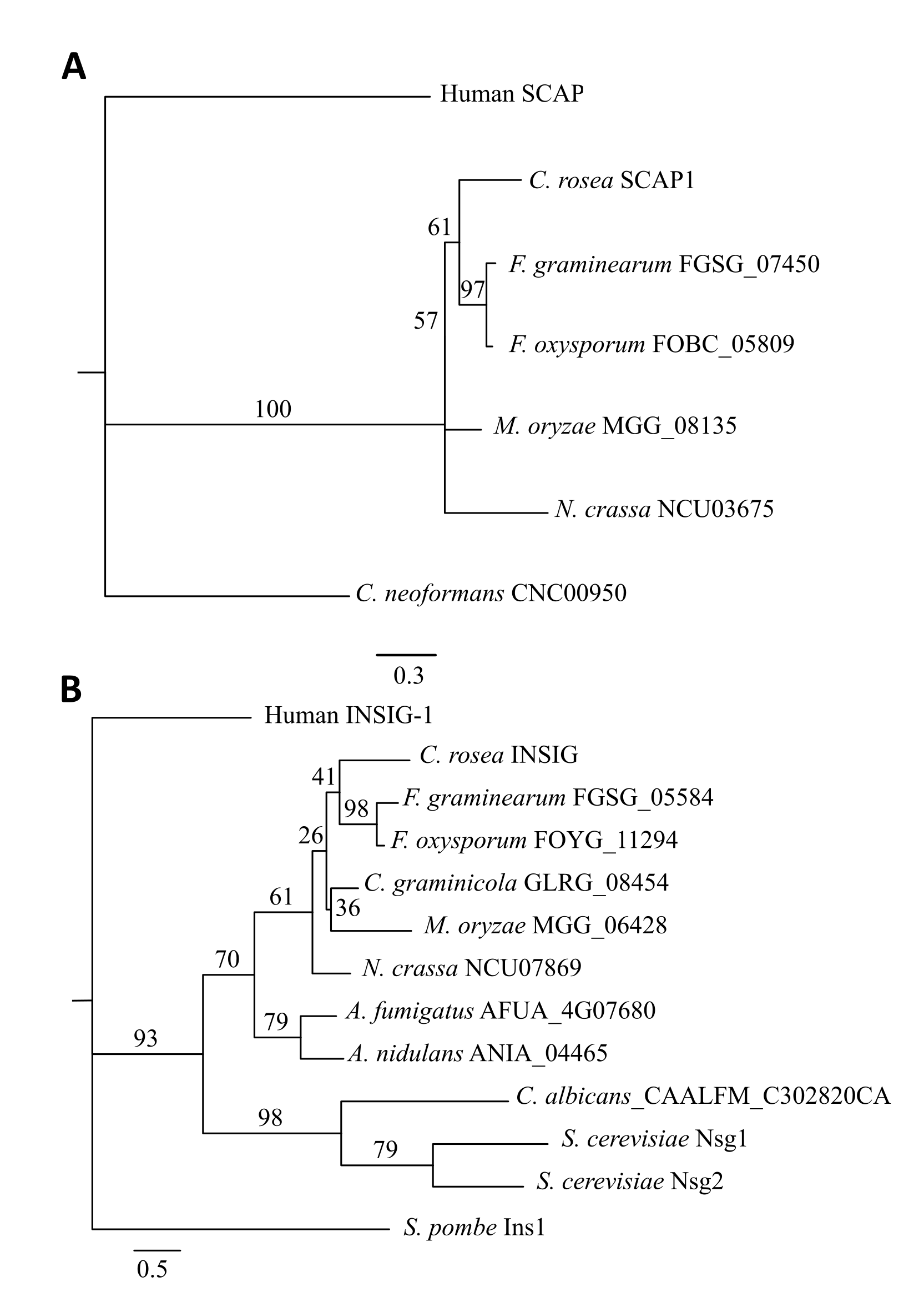

### Supplementary figure 4

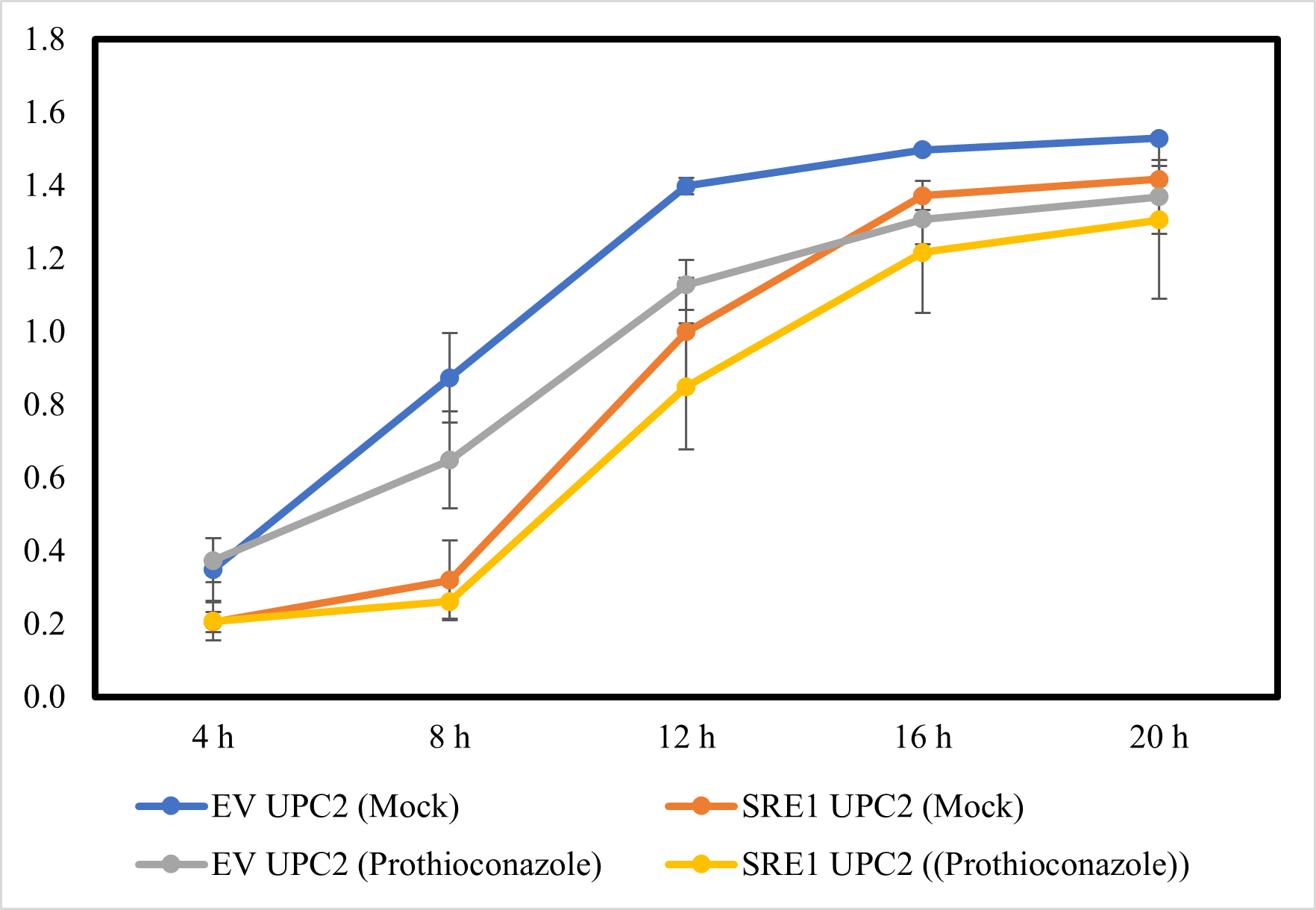

### Supplementary figure 5

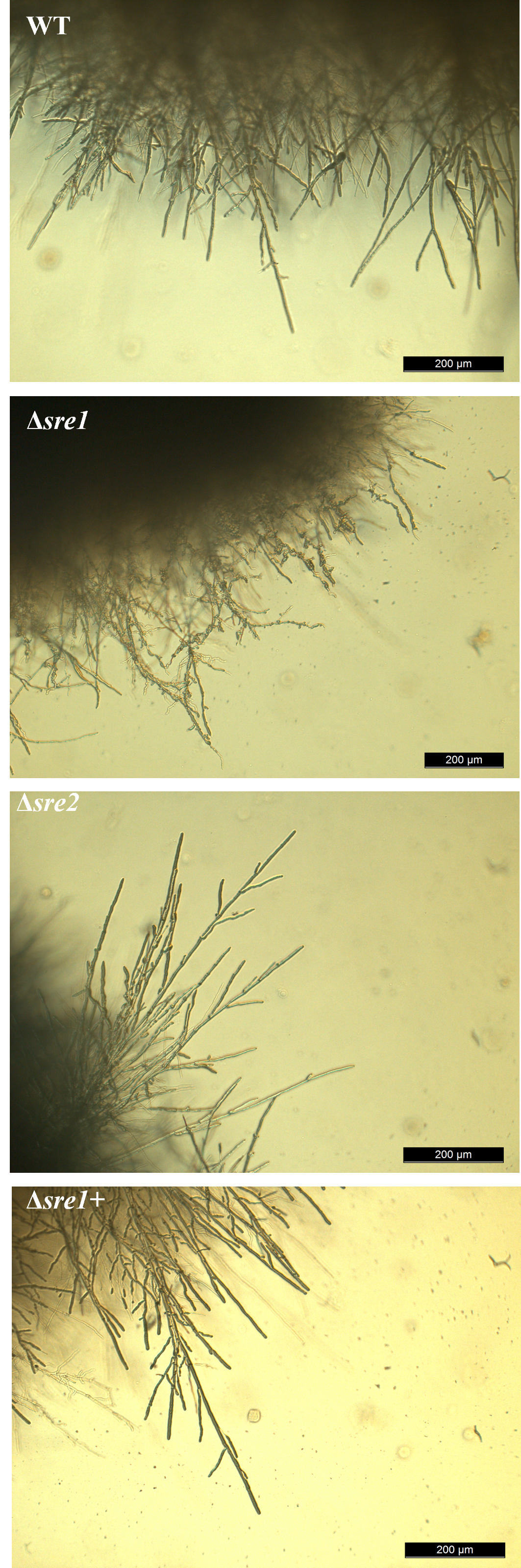

### Supplementary figure 6

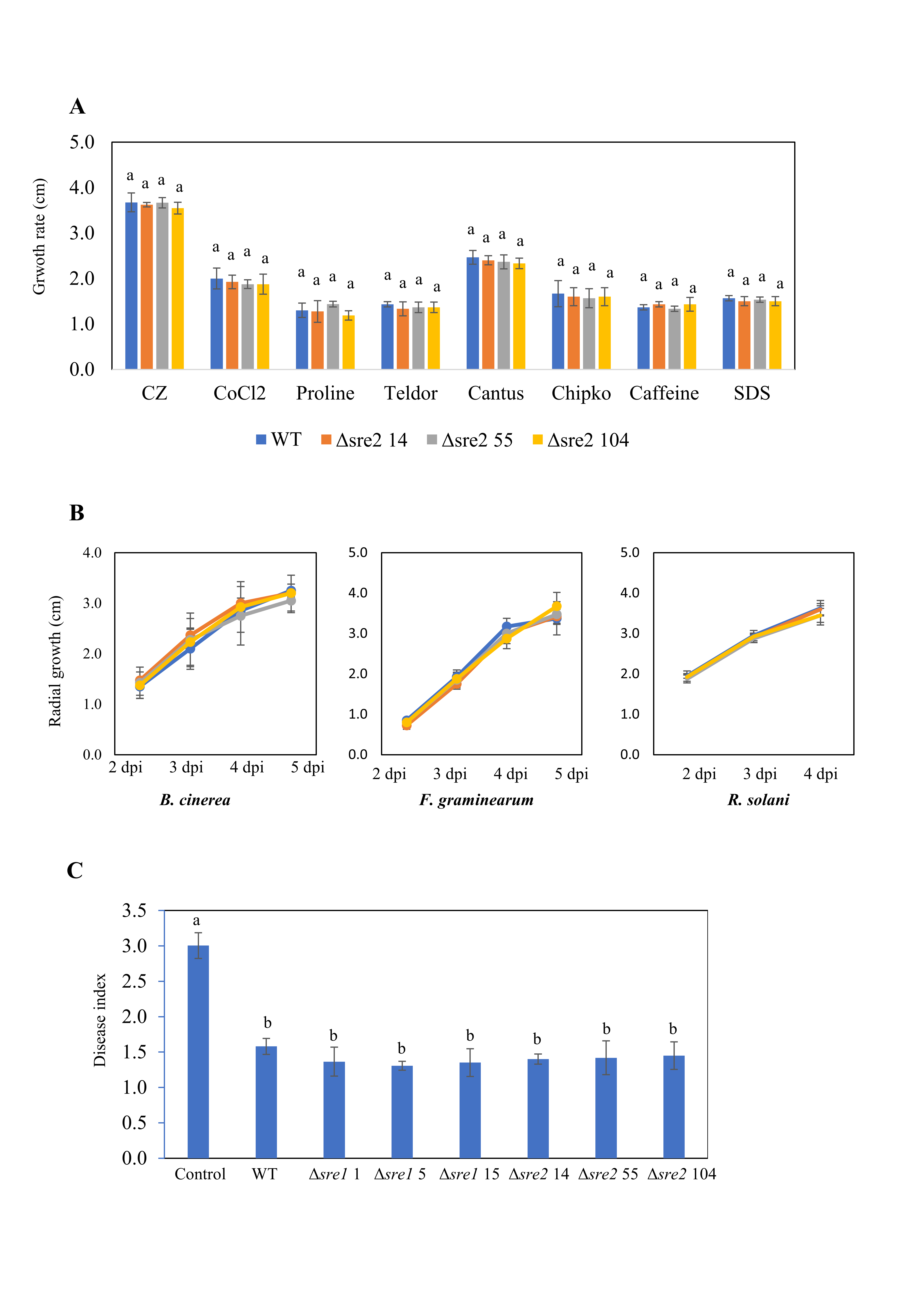
