## Supplementary table 2 for "SREBP-mediated gene expression regulation is essential for the intrinsic fungicide tolerance and antagonism in the fungal biocontrol agent *Clonostachys rosea*"

Table S2: List of primers used in this study.

| Primer name | 5’ to 3’ sequences | Use |
| --- | --- | --- |
| SRE1 ups F | ggggacaagtttgtacaaaaaagcaggcttatagataaggcccgcaggtgagt | To amplify *sre1* upstream |
| SRE1 ups R | ggggacaactttgtatagaaaagttgggtgtccggtgtcgagttgagtgtga |  |
| SRE1 ds F | ggggacaactttgtataataaagttgtaacctccagggccttcgtattgt | To amplify *sre1* downstream |
| SRE1 ds R | ggggaccactttgtacaagaaagctgggttgcgagtctggtttcacacgagt |  |
| SRE1 ko F | aggggccgtagaggtgggataata | *sre1* gene deletion validation |
| SRE1 ko R | ctaggggccaccaaccaaataagt |  |
| SRE1 qPCR F | gggacctcagcgaccacaagttca | *sre1* gene deletion validation |
| SRE1 qPCR R | cgaggttgtgggcacgctgaataa |  |
| SRE1 comp F | ggggacaagtttgtacaaaaaagcaggcttatagataaggcccgcaggtgagt | *sre1* complementation |
| SRE1 comp R | ggggacaacttttgtatacaaagttgtgcgagtctggtttcacacgagt |  |
| SRE2 ups F | ggggacaagtttgtacaaaaaagcaggcttatttttggttgcgtcatagcgtctg | To amplify *sre2* upstream |
| SRE2 ups R | ggggacaactttgtatagaaaagttgggtgtcatacggagacacggcgatttc |  |
| SRE2 ds F | ggggacaactttgtataataaagttgtaggtggccgggctggagat | To amplify *sre2* downstream |
| SRE2 ds R | ggggaccactttgtacaagaaagctgggtttggcgagctaatcttaaggcagtt |  |
| SRE2 ko F | caggagttggaggcggaaat | *sre2* gene deletion validation |
| SRE2 ko R | cgggcaaattccctttcgttagc |  |
| SRE2 qPCR F | ccaagcaggggcagcagttc | *sre2* gene deletion validation |
| SRE2 qPCR R | tccccctgacgacggttgct |  |
| Hyg F | gcgcgcaattaaccctcac | Mutant validation |
| Hyg R | gaattgcgcgtacagaactcc |  |

attB sequences for gateway cloning are underlined.
